# F26BP enables control of glycolysis rate independent of energy state

**DOI:** 10.64898/2026.01.31.703051

**Authors:** Megan M. Kober, Xinyi Yang, Bradley A. Webb, Denis V. Titov

## Abstract

Glycolysis is a conserved metabolic pathway that produces ATP and biosynthetic precursors. Multiple allosteric regulators control glycolytic enzymes *in vitro*. For example, phosphofructokinase (PFK) is allosterically regulated by fructose-2,6-bisphosphate (F26BP), ATP, ADP, AMP, citrate, acyl-CoA, and inorganic phosphate. It is not well understood which properties of homeostasis are enabled by each of these regulators, and whether they perform redundant or distinct functions. Using mathematical modeling and experiments with human cells lacking F26BP, we demonstrate that F26BP alters glycolytic rate independent of cellular ATP demand–a unique function not shared by other regulators. We also identified several downstream glycolytic intermediates as novel regulators of F26BP levels. Our findings clarify the role of F26BP as a unique regulator that controls the glycolytic rate independently of the cellular energy state in response to hormone and biosynthetic precursor levels. The F26BP regulatory circuit enables respiratory fuel selection and biosynthesis from glycolytic intermediates.

## INTRODUCTION

Glycolysis is a conserved metabolic pathway that produces both ATP and biosynthetic precursors. Extensive biochemical studies have established that four glycolytic enzymes are allosterically regulated by a dozen effectors, namely: hexokinase (HK), phosphofructokinase (PFK), glyceraldehyde-3-phosphate dehydrogenase (GAPDH), and pyruvate kinase (PK). However, it is largely unknown what physiological properties are enabled by allosteric regulation. We have recently demonstrated that allosteric regulation of HK and PFK by ATP, ADP, inorganic phosphate (P_i_), and glucose-6-phosphate (G6P) is required to maintain high ATP levels and prevent uncontrolled accumulation of phosphorylated glycolytic intermediates by inhibiting the Harden and Young reaction^1^. A surprising result of that study was that one of the classical regulators of glycolysis–fructose-2,6-bisphosphate–appeared to be dispensable for the regulation of glycolytic ATP production.

Fructose-2,6-bisphosphate (F26BP) is a potent allosteric activator of the glycolytic enzyme PFK and an allosteric inhibitor of gluconeogenic enzyme fructose-1,6-bisphosphatase (FBP)^2–4^. No other functions of F26BP are known, as it neither regulates nor participates in any other metabolic pathway. In mammals, F26BP levels are determined by a family of bifunctional enzymes, 6-phospho-2-fructokinase / fructose-2,6-bisphosphatase (PFKFB). Therein, the kinase domain synthesizes F26BP from ATP and glycolytic intermediate fructose-6-phosphate (F6P), while the phosphatase domain hydrolyzes F26BP to regenerate F6P^5–7^. In humans, four genes encode for PFKFB (PFKFB1; 2; 3; 4), each of which has distinct kinetic properties and exhibit tissue-specific expression^7–11^. It has also been proposed that TIGAR [TP53(tumor protein 53)-induced glycolysis and apoptosis regulator] is a fructose-2,6-bisphosphatase but other studies suggested that F26BP may not be its physiological substrate^12,13^.

PFKFB’s kinase and phosphatase activities are regulated by allosteric regulators and posttranslational modifications, allowing F26BP levels to change in response energy state, biosynthetic precursors, and hormone levels^14–24^. Genetic and pharmacological manipulation of PFKFB in mammalian cells alters glycolytic and gluconeogenic fluxes, confirming that F26BP is a physiologically important regulator of glycolytic and gluconeogenic rates in mammalian cells^25–33^. Yet, the knockout of all *S. cerevisiae* PFKFBs yielded no major phenotype besides changing the concentrations of PFK’s substrates and products, suggesting that other regulators may compensate for the lack of F26BP^34,35^. The complete knockout of all four mammalian PFKFB genes has not been investigated. Thus, the unique physiological function of F26BP remains incompletely understood, given its potential redundancy with other allosteric and hormonal regulators of PFK and FBP. For example, PFK’s allosteric activation by ADP and P_i_, FBP’s allosteric inhibition by AMP, and hormones that alter PFK and FBP activity through posttranslational modifications and gene expression all seemingly bypass F26BP’s allosteric effect.

Here, we aimed to investigate the unique advantages enabled by the F26BP regulatory circuit in human cells, using a combination of modeling and experiments. Our biophysical model of glycolysis predicted that F26BP enables cells to modulate glycolytic rate independent of ATP demand. Specifically, the model predicted that F26BP can increase the rate of glycolysis when respiration also generates ATP, but has no effect when glycolysis is the sole source of ATP. To validate this prediction, we generated human cell lines with *PFKFB1-4* quadruple knockout and an inducible F26BP-rescue system. By pointedly manipulating F26BP concentrations in human cells, we validated this prediction and further demonstrated that an F26BP-mediated increase in glycolytic rate occurs with a compensatory reduction in respiration rate. To better understand how cells control F26BP levels, we screened a library of core metabolic intermediates against PFKFB3 and identified both novel and strong inhibitors, including: 2-phosphoglycerate (2PG), phosphoenolpyruvate (PEP), cis-Aconitate, and citrate. Follow-up experiments with reconstituted glycolysis showed that PEP-mediated control of F26BP is important for matching glycolytic flux with biosynthetic demand. Collectively, these findings indicate that F26BP integrates signals from glycolytic intermediates and hormone levels, enabling cells to regulate glycolysis rate independently of energy demand to support biosynthesis and respiratory fuel selection.

## RESULTS

### Mathematical Model Predicts F26BP Uncouples Glycolytic Rate from Energy Demand

To investigate the function of F26BP allosteric regulation, we utilized our published mathematical model of mammalian glycolysis^1^. Briefly, this model utilizes enzyme rate equations derived from *in vitro* enzyme kinetics to simulate the activity of all glycolytic enzymes using a system of ordinary differential equations (ODEs) (Figure S1 A, B). Upon defining extracellular glucose concentrations and cellular ATP demand (ATPase), the model predicts time-dependent and steady-state concentrations of all glycolytic intermediates (Figure 1A). The published model contains a PFK rate equation that describes the response of PFK to F26BP based on extensive *in vitro* enzyme kinetic data, but does not yet account for the PFKFB enzymes that produce F26BP.

**Figure 1.**
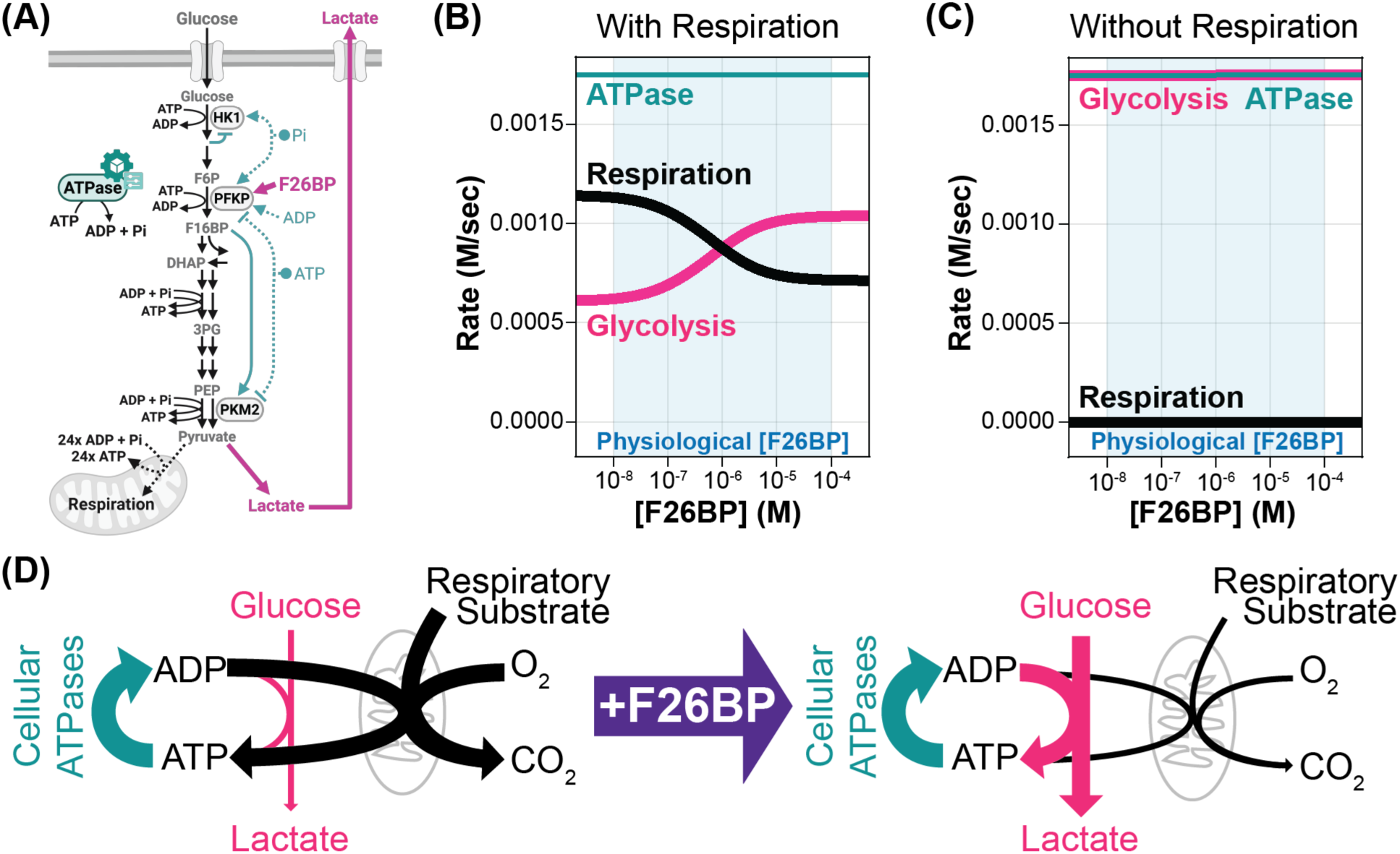
Model Predicts F26BP uncouples glycolytic rate from energy demand. (A) Schematic of glycolysis regulation accounted for in the biophysical model. Notable glycolytic intermediates (grey text), regulated enzymes (grey ovals), and allosteric regulators (pink or teal text) are highlighted. The model predicts that F26BP allosteric activation (solid pink lines) overrides glycolysis regulation by energy status (teal & black dashed lines). (B-C) Model simulations showing impact of F26BP on glycolytic (pink) and respiration (black) rates at a fixed ATPase reaction rate (teal) with (B) both glycolysis and respiration available to produce ATP and (C) only glycolysis available to produce ATP. The light blue shaded region indicates the physiological F26BP concentration range (10^-8^ to 10^-4^ M). (D) Schematic of F26BP’s predicted function in regulating glycolytic rate.

We simulated the effect of F26BP on glycolysis in two distinct metabolic states. The first is where glycolysis is the only ATP-producing pathway. The second state is where respiration, modelled as an ATP-producing reaction, also contributes to ATP production. We observed that F26BP increased the glycolytic rate when respiration produced ATP (Figure 1B), but had no effect when glycolysis produced all of ATP (Figure 1C). This aligns with our previous finding that the ability of glycolysis to match ATP supply with demand is driven by mass action and does not require allosteric regulation. The effect of F26BP on glycolytic rate was larger at low glycolytic rates than at high glycolytic rates (Figure S2 A-C), likely due to the fact that PFK is already partially activated at high glycolytic rates and thus cannot be activated by F26BP as effectively. Note that the total ATP production rate by glycolysis and respiration remains constant, so any increase in glycolytic ATP production rate by F26BP must yield a decrease in ATP production by respiration by the exact same amount. Thus, in cells, F26BP would be predicted to decrease respiration by the same amount as it activates glycolysis.

Collectively, these predictions suggest that the function of F26BP is to control glycolytic rate independent of ATP demand (Figure 1D). Such an ability could be central to supporting *de novo* biosynthesis from lower glycolytic intermediates or to enabling cellular fuel selection by prioritizing glucose over alternative respiratory fuels.

### Generation and validation of human cells devoid of F26BP

To test model predictions, we generated several cell lines that have variable levels of F26BP. First, we generated and validated the first reported human cell line that is completely devoid of F26BP. We eliminated F26BP from HeLa cells through iterative applications of CRISPR/Cas-9 and gRNAs targeting *PFKFB1, PFKFB2, PFKFB3,* and *PFKFB4* (see Table S1 for gRNA sequences; Figure S2 A-I). To enable rescue experiments, we introduced either Luciferase (negative control) or PFKFB3 with a C-terminal Flag tag under the control of a doxycycline-inducible promoter (TRE3G) into both wildtype HeLa (WT) and *PFKFB1-4* knockout (*PFKFB_0_*) cell lines. The *PFKFB_0_* and reintroduction of PFKFB3 were validated using sequencing, western blots, and intracellular F26BP measurements (Figure 2A, B; see Table S2 for sequencing primers). PFKFB3 overexpression in WT cells caused >10-fold increase in F26BP levels, consistent with the very large ratio of kinase to phosphatase activity of PFKFB3^36^. We observed no detectable changes in glycolysis or TCA cycle enzyme expression (Figure 2C, D), suggesting that any changes in the activity of these pathways would be driven by changes in enzyme activity.

**Figure 2.**
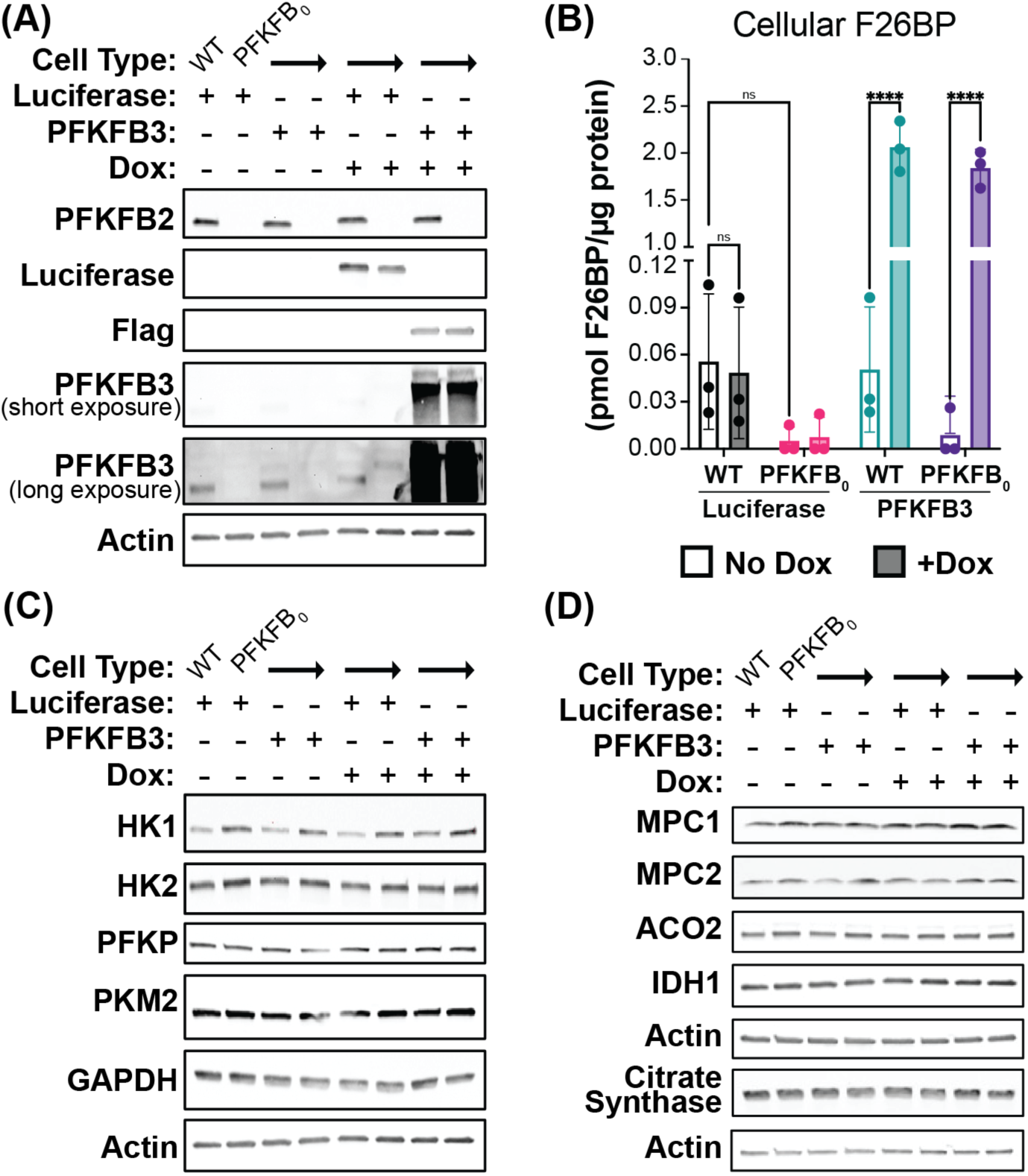
Generation and validation of PFKFB_0_ cells. **(A)** Western blot showing expression of Luciferase, PFKFB2, and PFKFB3 after 24-hour induction with water or 300 ng/ml doxycycline (Dox) in wildtype (WT) and quadruple *PFKFB1-4* knockout (PFKFB_0_) HeLa cells engineered to express Luciferase or PFKFB3-Flag with dox-inducible promoter. Endogenous PFKFB2 & PFKFB3 are visible in WT HeLa. Representative gel from one of three independent experiments. **(B)** F26BP levels in WT Luciferase (black), PFKFB_0_ Luciferase (pink), WT PFKFB3 (teal), & PFKFB_0_ PFKFB3 (purple) cell lines treated -/+ Dox for 24 hours (- /+ shading, respectively). One hour prior to F26BP extraction, cells were treated with a vehicle control (DMSO). F26BP extracts were normalized to cell protein. Mean ± SD from n=3 independent experiments. P-values were calculated using 2-way ANOVA followed by Tukey’s multiple comparisons test. ns p > 0.05, ****p < 0.0001. **(C)** Western blot of glycolytic enzymes and **(D)** TCA cycle enzymes in WT Luciferase, PFKFB_0_ Luciferase, WT PFKFB3-Flag, & PFKFB_0_ PFKFB3-Flag cell lines treated -/+ Dox for 24 hours. Representative gel from one of two independent experiments.

### F26BP increases glycolytic rate and decreases respiration rate in human cells

With the ability to experimentally manipulate cellular F26BP, we wanted to test the model’s prediction that F26BP increases glycolytic rate while decreasing respiration rate without a net change in ATP production rate (see Figure 1B for model predictions). In WT cell lines, glycolytic fermentation contributes to ∼70% of the total ATP production rate compared to respiration (Figure 3A). Despite basal glycolytic rates operating near maximal rates, PFKFB3 overexpression in both WT and *PFKFB_0_* cell lines doubled the ratio of glycolysis to respiration (Figure 3B, C). In complete agreement with model predictions, PFKFB3 overexpression prompts a small increase in glycolysis and large inhibition of respiration (Figure 3C, D; Figure 2S A-C). In alignment with this observation, the *PFKFB_0_* cells with and without Luciferase expression exhibit slightly reduced glycolytic rates and elevated respiratory rates compared to WT (Figure 3C, D). Collectively, these observations fully recapitulate the predictions of our model that F26BP increases the glycolytic rate with a compensatory reduction in the respiration rate without a change in net ATP production rate.

**Figure 3.**
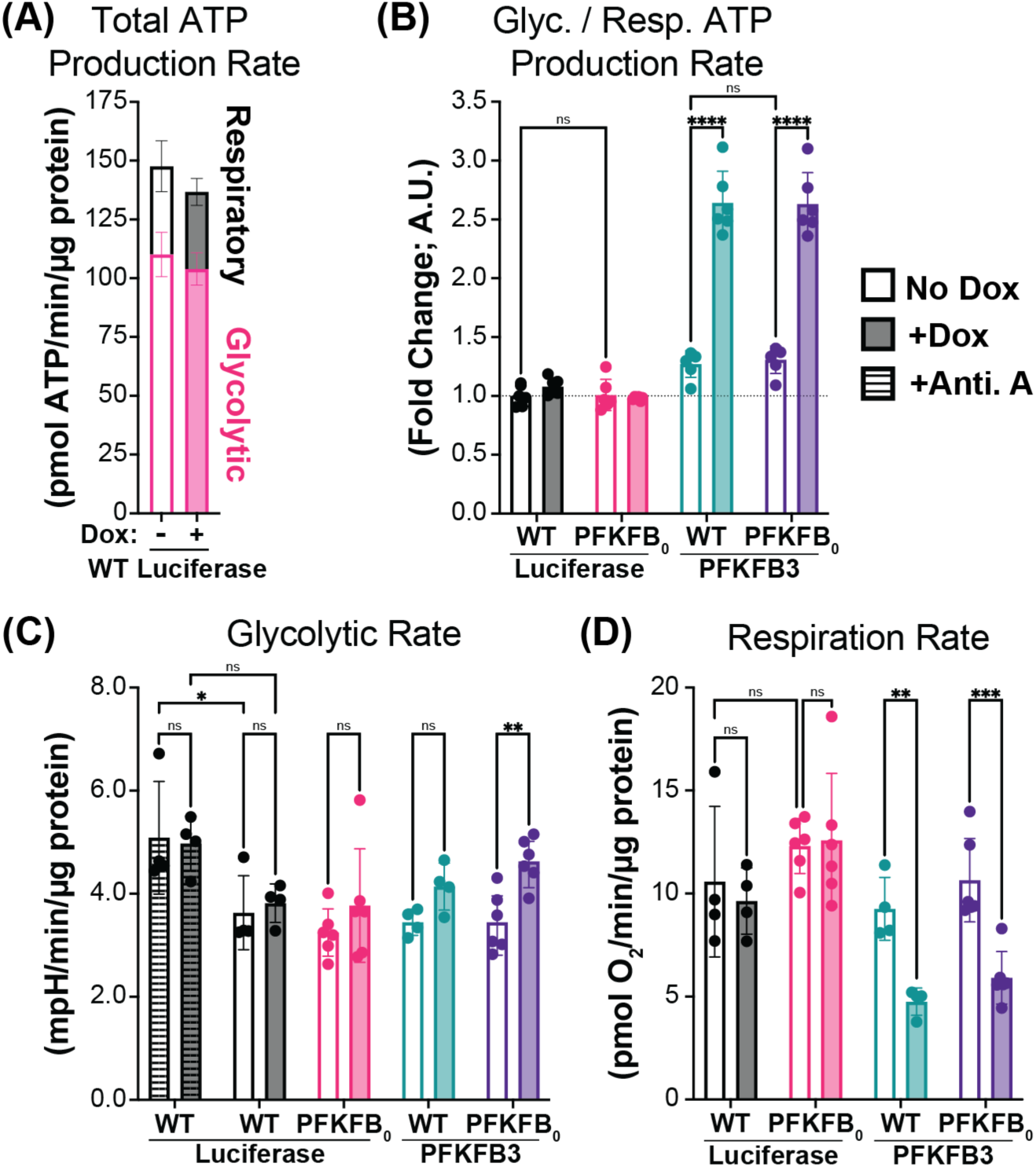
Cellular F26BP determines glycolytic rate in the presence of respiration. (A) The contributions of glycolysis (pink) and respiration (black) to the total ATP production rate in WT Luciferase cells -/+ Dox for 24 hours (-/+ shading, respectively). Mean ± SD from n=2 independent experiments. (B) PFKFB3 expression increases the relative glycolytic to respiration ATP production rate ratio. WT Luciferase (black), PFKFB_0_ Luciferase (pink), WT PFKFB3 (teal), & PFKFB_0_ PFKFB3 (purple) cell lines treated -/+ Dox for 24 hours (-/+ shading, respectively). Presented as a fold-change relative to WT Luciferase without Dox treatment. Mean ± SD from n=2 independent experiments. (C) Effect of Luciferase and PFKFB3 expression on basal (no stripes) and maximal (stripes) glycolytic rates. Treatment with Antimycin A (Anti. A) achieved maximal glycolytic rates and serves as a positive control. Quantified by extracellular acidification rates, normalized to cell protein. Mean ± SD from n=2 independent experiments. (D) Effect of Luciferase and PFKFB3 expression on basal respiration rates. Quantified by oxygen consumption rates, normalized to cell protein. Mean ± SD from n=2 independent experiments. P-values were calculated using 2-way ANOVA followed by Tukey’s multiple comparisons test. ns p > 0.05, ** p < 0.01, *** p < 0.001, **** p < 0.0001.

### F26BP does not affect glycolytic rate when respiration is inhibited

We next tested the model prediction that F26BP does not alter glycolytic rates in the absence of respiration. To inhibit respiration and elicit maximal glycolytic rates, we treated cells with electron transport chain complex III inhibitor, Antimycin A (Figure 4A). We then confirmed that Antimycin A did not significantly change F26BP concentrations in WT and *PFKFB_0_* in the presence or absence of PFKFB3 overexpression (Figure 4B). Despite all experimental manipulations to F26BP, we observed no significant difference in the maximal glycolytic rates achieved by WT and *PFKFB_0_* cell lines with or without PFKFB3 overexpression (Figure 4C). Together, these observations validate the model prediction that allosteric regulation by F26BP does not affect glycolytic rates in the absence of respiration.

**Figure 4.**
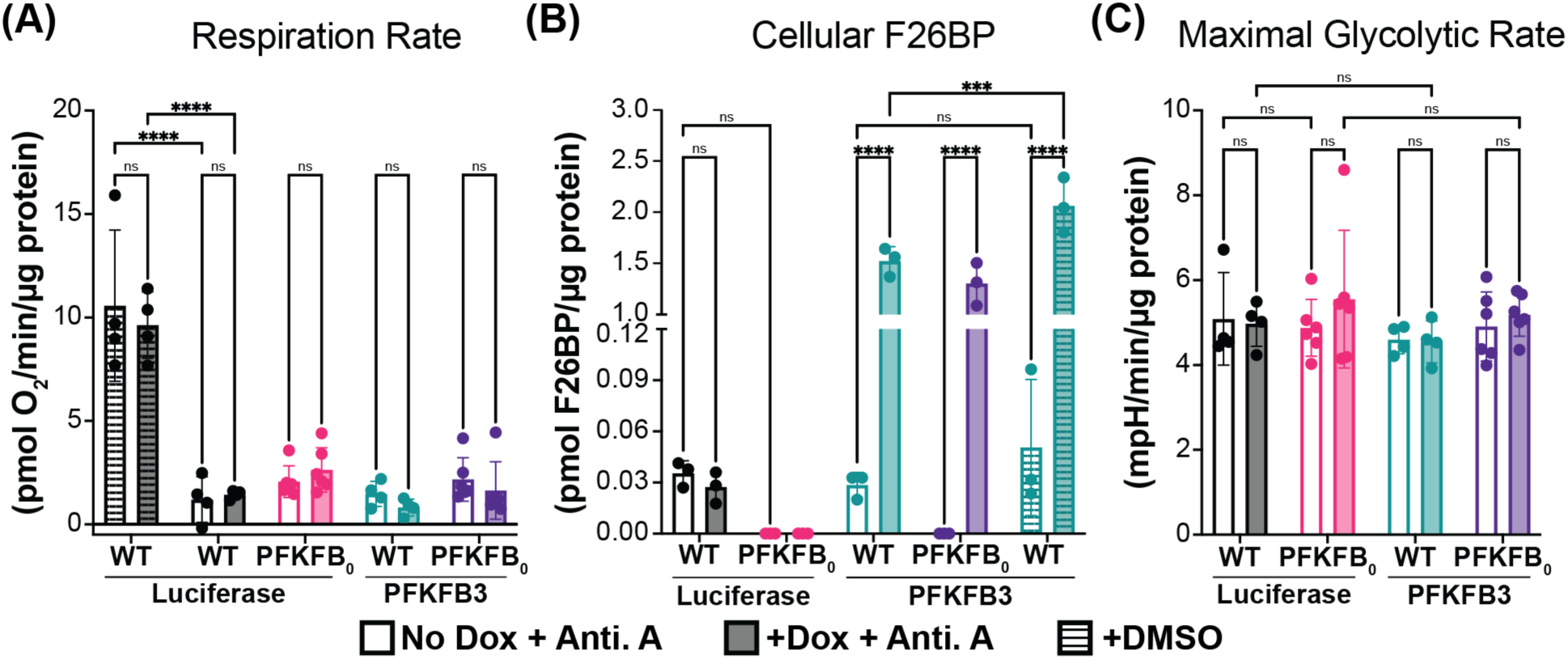
Cellular F26BP does not impact maximal glycolytic rates in the absence of respiration. (A) Basal (stripes) and inhibited (no stripes) respiration rates in WT Luciferase (black), PFKFB_0_ Luciferase (pink), WT PFKFB3 (teal), & PFKFB_0_ PFKFB3 (purple) cell lines treated -/+ Dox for 24 hours (-/+ shading, respectively). Respiration rates were inhibited using Antimycin A (Anti. A) treatment, quantified by oxygen consumption rates, and normalized to cell protein. Mean ± SD from n=2 independent experiments. (B) Effect of Anti. A treatment on F26BP levels. One hour prior to F26BP extraction, cells were treated with Anti. A (no stripes) or a vehicle control (DMSO; stripes). F26BP extracts were normalized to cell protein. Mean ± SD from n=3 independent experiments. (C) Effect of Luciferase and PFKFB3 expression on maximal (stripes) glycolytic rates. Maximal glycolytic rates were achieved using Anti. A treatment, quantified by extracellular acidification rates, and normalized to cell protein. Mean ± SD from n=2 independent experiments. (P-values were calculated using 2-way ANOVA followed by Tukey’s multiple comparisons test. ns p > 0.05, ** p < 0.01, *** p < 0.001, **** p < 0.0001.

### *In vitro* screen of PFKFB activity identifies novel regulators of F26BP levels

Having shown that F26BP controls the rate of glycolysis in the presence of respiratory ATP production, we wondered how intracellular metabolites regulate F26BP itself. Because PFKFB activity is the sole determinant of F26BP concentrations in human cells (Figure 5A), we conducted an *in vitro* regulator screen using recombinant PFKFB3 (Figure 5B) and quantifying F26BP production (Figure S4 A-D). We prioritized characterizing PFKFB3 due to its high proteomic abundance in proliferating cells, thereby ensuring compatibility with the mathematical glycolysis model and cell systems utilized in this study^37–39^. Metabolites included in the screen spanned major roles and destinations of glycolysis (i.e., energy and redox maintenance, the TCA cycle, *de novo* amino acid and lipid synthesis), as well as known PFKFB3 regulators as controls (Figure S5 A-D; see Table S3 for complete list)^14–19^. PFKFB3 activity was evaluated under physiologically relevant conditions by quantifying F26BP in the presence and absence of the screened regulators (Figure 5C). As expected, increasing the substrate F6P and adding the known allosteric activator inorganic phosphate (P_i_) increased F26BP production up to 2-fold. While previously established allosteric inhibitors ADP, phosphoenolpyruvate (PEP), and citrate decreased F26BP production by at least half.

**Figure 5.**
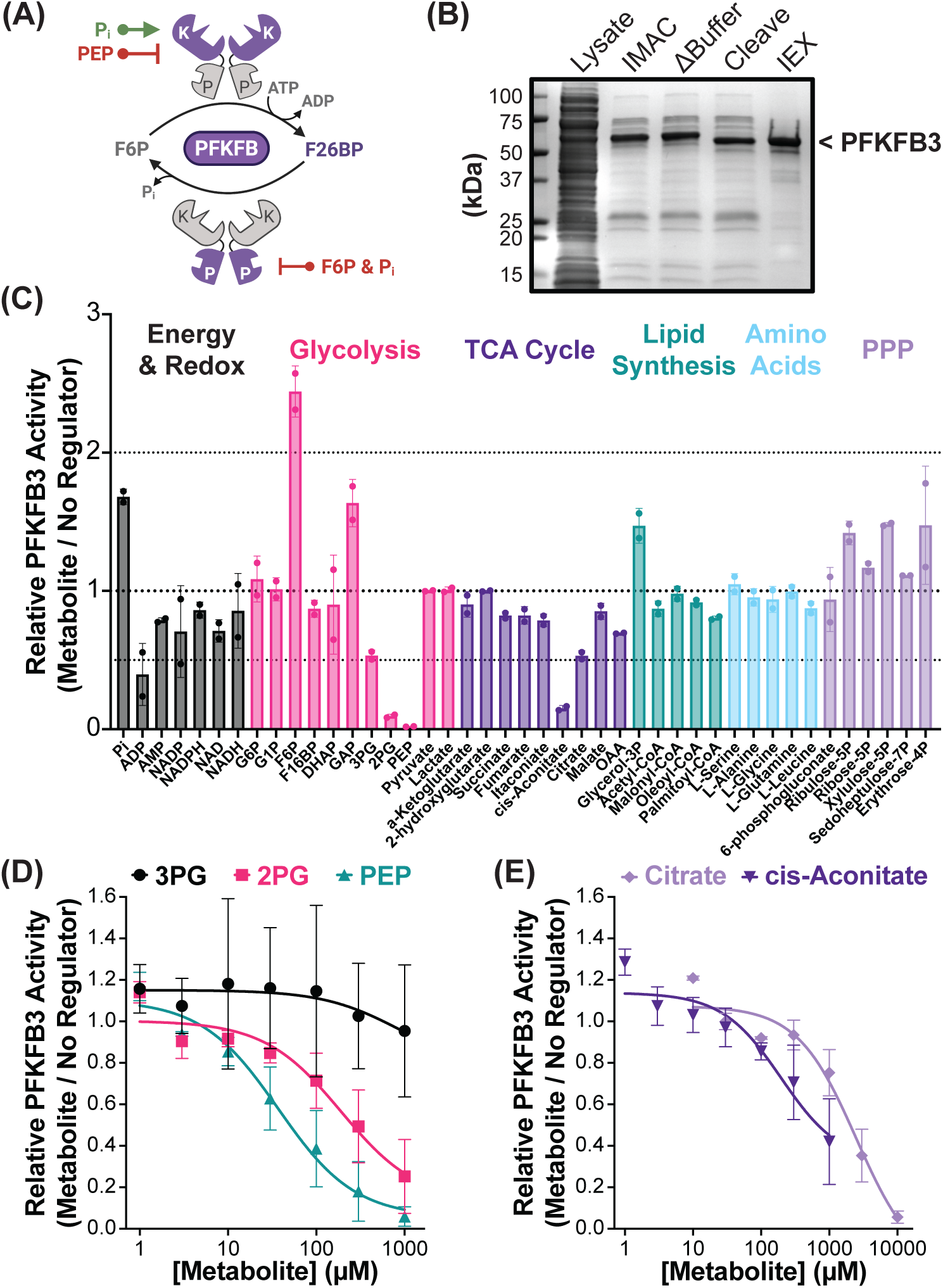
PFKFB activity screen identifies glycolytic & TCA Cycle intermediates as novel regulators of F26BP. (A) Schematic of 6-phosphofructo-2-kinase/ fructose-2,6-bisphosphatase (PFKFB) activity and previously established allosteric regulators (activators in green, inhibitors in red). Purple shading denotes kinase (K) or phosphatase (P) domain activity. (B) Coomassie showing recombinant PFKFB3 purification. His-PFKFB3 was isolated from bacterial lysate (Lysate) using immobilized metal ion affinity chromatography (IMAC). Protease cleavage (Cleave) removed His tag prior to purification via anion exchange chromatography (IEX). Representative gel from one of two independent purifications. (C) Relative PFKFB3 activity in the presence of physiological substrates (0.2 mM F6P and 3 mM ATP) and metabolites involved in: energy & redox balance (black), glycolysis (pink), the TCA Cycle (purple), lipid synthesis (teal), non-essential amino acid synthesis (blue), & Pentose Phosphate Pathway (PPP; lavender). All regulators had a final concentration of 1 mM, except Acetyl-CoA, Malonyl-CoA, Oleoyl-CoA and Palmityl-CoA at 0.1 mM. Presented as fold-change relative to no regulator control. Mean ± SD from n=2 independent experiments. (D-E) Relative PFKFB3 activity in the presence of identified inhibitor concentrations with best-fit agonist response curve (solid line; Hill’s Coefficient = 1). Presented as fold-change relative to no regulator control. Mean ± SD from n=3 independent experiments. (D) Titration of inhibitory glycolytic intermediates: 3-phosphoglycerate (3PG; black circles), 2-phosphoglycerate (2PG; pink up triangles), & phosphoenolpyruvate (PEP; teal down triangles). (E) Titration of inhibitory TCA Cycle intermediates: cis-Aconitate (purple diamonds) & citrate (lavender squares).

Most notably our screen identified several novel inhibitors of human PFKFB3, including: the lower glycolytic intermediates 3-phosphoglycerate (3PG) and 2-phosphoglycerate (2PG), as well as the citrate-derivative cis-Aconitate. In further examining PFKFB3’s inhibition by the lower glycolytic intermediates (Figure 5D), we found PEP was the most potent effector. Although both 2PG and 3PG were less potent compared to PEP, we found that 2PG was still able to inhibit PFKFB3 at physiologically relevant concentrations (10-100 μM). Similarly, in examining the TCA cycle derivatives (Figure 5E), both citrate and cis-Aconitate were able to inhibit PFKFB3 within physiological ranges (5 mM and 1 mM, respectively)^40^.

### PFKFB enables the regulation of glycolytic flux by PEP levels

The regulation of F26BP levels by PEP is functionally analogous to a feedback inhibition of PFK activity by PEP, which is the mode of regulation observed in bacterial but not eukaryotic PFK^41^. To examine how PFKFB3 regulation could enable pathway feedback inhibition by lower glycolytic intermediates within mammalian glycolysis, we utilized a partial reconstitution of glycolysis (Figure 6A) that measures flux through the PFK platelet isoform (PFKP). PFKP activity was evaluated in both the presence and absence of PFKFB3, and therefore with and without regulated F26BP production. The addition of PFKFB3 increases PFKP activity 10-fold due to the potent allosteric activation by F26BP (Figure 6B). Therein, it is apparent that PFKP is markedly inhibited under a physiologically relevant ATP/ADP ratio (10:1) due to its severe allosteric inhibition by ATP, as has been previously suggested^42^. Notably, we showed that PEP is only able to inhibit glycolysis flux in the presence of PFKFB3, validating that PEP enables regulation of glycolysis by lower glycolysis intermediates (Figure 6B). We further examined the relevance of PFKFB3’s inhibition by 2PG and similarly demonstrated an opportunity pathway feedback inhibition – albeit with less potency compared to PEP (Figure 6C).

**Figure 6.**
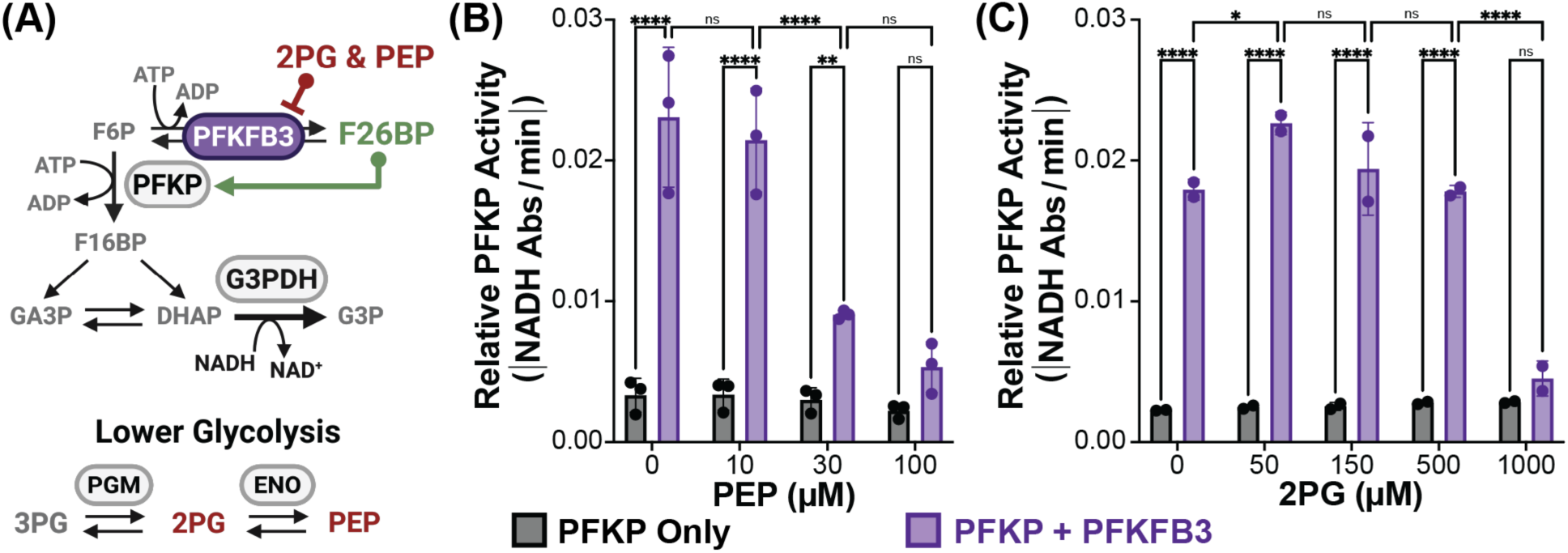
Regulation of PFKFB3 controls flux through rate-limiting glycolytic enzyme PFKP. (A) Schematic of F26BP regulatory circuit. Notable intermediates (grey text), enzymes (grey ovals, except PFKFB3 in purple), and allosteric regulators (red or green text) are highlighted. *In vitro* upper glycolysis assay terminates with glycerol-3-phosphate dehydrogenase (G3PDH), which enables continuous reaction monitoring by NADH consumption (black text). (B-C) PFKP activity in the absence (black) or presence of PFKFB3 (purple) with increasing lower glycolysis intermediate concentrations. F26BP regulatory circuit with: (B) Phosphoenolpyruvate (PEP) titration, mean ± SD from n=3 independent experiments, and (C) 2-phosphoglycerate (2PG) titration, mean ± SD from n=2 independent experiments. P-values were calculated using 2-way ANOVA followed by Tukey’s multiple comparisons test. ns p > 0.05, * p < 0.05, ** p < 0.01, *** p < 0.001, **** p < 0.0001.

## DISCUSSION

Here, we investigated the function of the PFK activator, F26BP, in the regulation of glycolysis. Specifically, we focused on identifying a unique regulatory function that is enabled by the presence of the F26BP regulatory circuit, which cannot be performed by other regulators of glycolytic enzymes. Using a combination of modeling and experiments, we showed that F26BP increases glycolytic rate with a compensatory reduction in respiration rate. Our results also explain why experiments with PFKFB knockouts in *S. cerevisiae* showed no obvious phenotype, as respiration has not been measured in those studies^34,35^. We predict that the main effect of F26BP in highly glycolytic *S. cerevisiae* cells will be to inhibit respiration, as we observed for HeLa cells. Additionally, we showed that F26BP does not affect the total ATP production rate by the cell, and it has no effect on glycolysis rate in the absence of respiration. These findings are perhaps expected, as the ATP production rate is set by ATP demand and cannot be affected by the regulation of ATP-producing pathways. For example, if ATP production exceeds demand this will lead to a rapid fall in ADP levels. Because ADP is a necessary substrate for all ATP-producing pathways, ATP production rates will expectantly decrease and restore the balance between ATP production and consumption^1^.

A unique property of F26BP is that it is a purely regulatory metabolite that does not participate in other metabolic pathways. This allows cells to change the level of F26BP and thus glycolysis rate without directly affecting other parts of metabolism. Changes in F26BP levels will, of course, affect other metabolic pathways like respiration, but those are compensatory changes in response to glycolytic rate changes and not the direct effect of F26BP. Other allosteric regulators of glycolytic enzymes, such as ATP, ADP, inorganic phosphate, citrate, or acyl-CoA, are substrates and products in other metabolic pathways – making it impossible change their levels without directly impacting affiliated pathways^43–48^. This metabolic isolation of F26BP is likely an important factor in the evolution of the F26BP regulatory circuit, as it enables glycolytic rates to respond to hormonal and allosteric regulators independent of cellular energy status.

Notably, the regulation of PFK by F26BP appears to be a eukaryote-specific mechanism^49^. For example, *E. coli* PFK is inhibited by PEP directly, while human PFK is inhibited by PEP indirectly through inhibition of F26BP production by PEP^15–17,41^. While both mechanisms enable inhibition of PFK by PEP, we speculate that the presence of the F26BP circuit enables more versatile regulation of glycolytic rate in eukaryotes by allowing more regulators to affect PFK through a combination of direct binding to PFK and regulation of F26BP levels via direct binding to PFKFB. This idea is supported by the observation that human PFK is twice the size of *E. coli* PFK, due to the fusion of two bacteria-like PFK sequences and the mutation of one of the catalytic sites to an allosteric site, resulting in an increase of allosteric sites from one to three^50,51^. In humans, the combination of three PFK and four PFKFB genes with different kinetic properties and tissue-specific expression likely enables fine-tuning of glycolysis regulation that may be difficult to achieve using the same number of bacterial PFK genes.

To this end, mammals have evolved genetic, hormonal, and allosteric mechanisms to control F26BP levels, which enables glycolytic rates to be regulated in a tissue-specific manner. Such regulation of F26BP allows tissues to modify glucose utilization based on environmental factors. Specific examples include: downregulation of liver F26BP levels by glucagon in the fasted state to switch from glucose utilization to production, upregulation of muscle F26BP levels by insulin in the fed state to stimulate glucose oxidation, and upregulation of kinase dominant PFKFB3 isoform to stimulate fermentation in hypoxia and rapidly proliferating cells^7,10,24,38,39^. Furthermore, F26BP levels are regulated in such a way to increase glycolytic rates and support glucose-derived biosynthesis. For instance, inhibition of F26BP production by PEP and citrate provide feedback from biosynthetic precursors within glycolysis and the TCA cycle, while stimulation of F26BP production by insulin favors glucose-derived lipid synthesis in adipose^15–18,27,52,53^. Together with our findings, it is clear that the complexity of regulating F26BP has diverse metabolic implications, including enabling cellular fuel selection and supporting biosynthesis.

Finally, we highlight that the combined use of simulations and experiments was instrumental for the investigation of the function of F26BP reported here. We believe that simulations are an indispensable tool for understanding the regulation of metabolic pathways. Most metabolic pathways have more than one allosteric enzyme with more than one effector, which makes it nearly impossible to understand their combined action without simulations. With the continued use of accurate *in vitro* enzyme kinetics equations, extensions of our glycolysis model to other pathways will make it feasible to simulate the activity of many metabolic pathways in human cells. By combining these endeavors with purposefully designed experiments, there is incredible potential to improve our understanding of metabolism to the point of accurately predicting metabolic states across different human cell types under variable conditions.

## METHODS

### Glycolysis Model Code

Simulations of glycolysis in the presence and absence of respiration were performed using an extension of our previously published glycolysis model implemented in CellMetabolism.jl^1^. The glycolysis model contains enzyme rate equations for all glycolytic enzymes derived from *in vitro* enzyme kinetics, including a Monod-Wyman-Changeux (MWC) rate equation for phosphofructokinase (PFK) that describes its allosteric regulation by F26BP, ATP, ADP, AMP, citrate, and inorganic phosphate. To model respiration, we added an ATP Synthase reaction that produces ATP from ADP and inorganic phosphate. The ATP Synthase rate equation follows reversible Michaelis-Menten kinetics with competitive product inhibition:

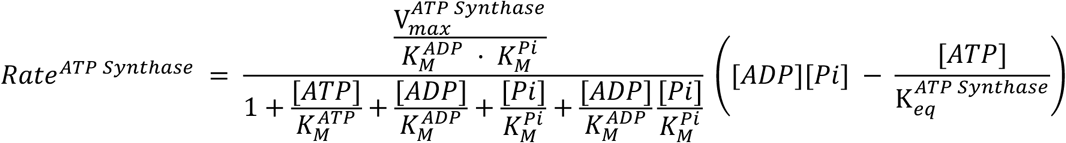

where 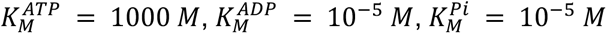, and *K_eq_* = 10^6^. Rate equation for ATP synthase was added to corresponding ordinary differential equations for ATP, ADP and Phosphate, which is done automatically by CellMetabolism.jl package. The system of ordinary differential equations was numerically simulated to steady state using the RadauIIA9 implicit Runge-Kutta solver from DifferentialEquations.jl package^54^ with absolute tolerance of 10^-15^ and relative tolerance of 10^-8^. For simulations varying F26BP concentration, F26BP was held constant at each specified level (range: 10^-8^ to 10^-4^) while all other metabolite concentrations were allowed to reach steady state. The respiration-to-glycolysis ATP production ratio was varied by adjusting 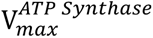 and glycolytic enzyme concentrations while keeping the sum of *V_max_* of glycolysis and respiration constant. Total ATP demand 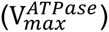 was kept at 20% of the sum of *V_max_* of glycolysis and respiration. Code to reproduce Figure 1B, C and Figure S2 is available at: https://github.com/DenisTitovLab/CellMetabolism.jl/tree/main/examples/2026_F26BP_Figures

### Cell Lines

HeLa cells were purchased from ATCC (CCL-2) and cultured in Dulbecco’s modified eagle’s medium [DMEM (Gibco™ #11995, see Table S4 for media composition), 10% fetal bovine serum (FBS) (Avantor #89510-186), and 100 U/mL penicillin–streptomycin (Gibco™ #15140122)] (cDMEM) and maintained in a 37°C in 5% CO_2_ incubator. Lentiviral-infected cells were cultured in DMEM (Gibco™ #11995), 10% FBS (Avantor #89510-186), 100 U/mL penicillin–streptomycin (Gibco™ #15140122), and 10 μg/mL Puromycin (Thermo Scientific #A1113803) and maintained in a 37°C in 5% CO_2_ incubator. All experiments were conducted in the absence of penicillin, streptomycin, and puromycin. Cells were regularly tested for mycoplasma and were mycoplasma free.

### Generation of Quadrupole Knock-Out Cells via Transient CRISPR/Cas9

The *PFKFB_0_* cell lines were generated by conducting 4 rounds of a high deletion frequency CRISPR/Cas9 protocol previously described, wherein each round was defined by the PFKFB gene (*PFKFB1, PFKFB2, PFKFB3,* or *PFKFB4)* being targeted^55,56^. Two guide RNAs (gRNA-A and gRNA-B) were designed per PFKFB gene (Table S1) and subcloned into pSpCas9(BB) (pX459; Addgene #48139) using Golden Gate assembly (T4 PNK New England Biolabs #M0201, BbsI New England Biolabs #R3539, and T4 DNA Ligase New England Biolabs #M0202T).

Initially, 0.125 x 10^6^ HeLa(WT) were seeded in 6 cm dishes. Twenty-four hours after seeding, culture media was reduced to 2 mL and cells were transfected by adding 200 µL of transfection mixture. The transfection mixture contained 6 µL X-treme Gene 9 reagent (Roche #06365787001), 1 µg pX459-PFKFB1-gRNA-A, 1 µg pX459-PFKFB1-gRNA-B, and Opti-MEM (Life Technologies #31985-070) up to 200 µL. The mixture was incubated at room temperature for 30 minutes before adding dropwise to cells. Twenty-four hours after transfection, an additional 2 mL of cDMEM was added to the plate (total volume: ∼4 mL). Forty-eight hours post-transfection, cells were positively selected by treatment with cDMEM + 1.25 μg/mL puromycin and monitored daily. After all non-transfected cells died (typically, 48 hours post-selection), positively-selected cells were returned to cDMEM for the remainder of the validation and experimental protocols. Twenty-four hours after media exchange, individual cells were isolated from the surviving bulk culture by using a limiting dilution of ∼2 cells/200 µL and seeded in 96-well plates.

Once single-cell derived colonies reached a sufficient culture size (typically 2-3 weeks), knock out efficiency was screened by extracting genomic DNA using QuickExtract (Lucigen #QE0905T) and conducting non-deletion/deletion PCR as described (Table S2)^55,56^. Promising knock out clones were validated by on PCR amplified gRNA cut sites (Table S2) and analyzing the Sanger Sequencing results using the following methods: Tracking of Indels by DEcomposition (TIDE) and EditCo’s Inference of CRISPR Edits (ICE)^57–59^. Knock outs of endogenously expressed PFKFB2 and PFKFB3 were also confirmed by Western Blot. Once validated, multiple, genetically-distinct knock out clones were carried forward in iterative applications of this protocol targeting *PFKFB2*, *PFKFB3*, and *PFKFB4*, respectively.

### Generation of Stable Expressing Cells using Lentivirus

Lentivirus was generated by seeding 0.675 x 10^6^ HEK293T cells in a 6-well plate with 2 mL of cDMEM per well. Twenty-four hours after seeding, cells were transfected by adding 100 µL of transfection mixture per well. The transfection mixture contained 3 µL X-treme Gene 9 reagent (Roche #06365787001), 500 ng psPAX2 6 (Addgene #12260), 50 ng pMD2.G (Addgene #12259), 500 ng of the pLVX-TRE3G expression vector (Takara #631187, containing either human PFKFB3 with a C-terminal Flag-tag, or Luciferase as a control) and Opti-MEM (Life Technologies #31985-070) up to 100 µL. The mixture was incubated at room temperature for 30 minutes before adding dropwise to cells. Twenty-four hours post transfection, an additional 1 mL of cDMEM was added to each well (total volume per well: ∼3 mL). Forty-eight hours post transfection, lentivirus was harvested by collecting spent media from the transfected HEK293T, processed through a 0.45 um filter, and then immediately utilized to infect destination cell lines.

The destination cell lines were prepared for infection by seeding 75 x 10^3^ of the respective cell line (HeLa(WT) or *PFKFB_0_*) per well of a 6-well plate in 2 mL of cDMEM. Twenty-four hours after seeding, the media of destination cell line media was reduced to 1 mL per well. Cells were then infected by adding 2 mL of an infection solution to each well (final volume per well: 3 mL). The infection solution consisted of 12 μg/mL polybrene (final: 8 μg/mL) and fresh lentivirus titration (final: 2000-50 μL) diluted in cDMEM. Twenty-four hours post-infection, the media was replaced with fresh cDMEM. Forty-eight hours post-infection, stable cell lines underwent positive selection for at least a week by exchanging media to cDMEM + 1 μg/mL Puromycin (Thermo Scientific #A1113803). For ongoing maintenance, high expression cell lines were cultured in 10 μg/mL Puromycin (Thermo Scientific #A1113803).

### SDS-PAGE and Western Blotting

0.2 x 10^6^ cells were seeded per well of 6-well plate in 2 mL of cDMEM. Twenty-four hours after seeding, 2 mL of cDMEM was added per well supplemented with doxycycline (final: 300 ng/mL; Sigma #D9891) or water. Twenty-four hours after induction, protein lysates were extracted by treating with an SDS lysis buffer (50 mM Tris (pH 8), 1 mM EDTA, 1% SDS, and cOmplete Mini EDTA-free Protease Inhibitor Cocktail (Roche #11836170001)) and boiling at 95°C for 10 minutes. Cellular protein was measured by Pierce™ bicinchoninic acid (BCA) protein assay kit (Thermo Fisher Scientific #23227) to equalize protein quantities across samples. 10-20 μg of total cellular protein was analyzed by mixing with Laemmli buffer (BioRad #1610747, prepared in accordance to manufacturer’s instructions), separating by gel electrophoresis on 4–20% gradient Tris-Glycine gels (BioRad Mini-Protean TGX #4561094), and transferring onto nitrocellulose membranes using the BioRad Trans-Blot Turbo Transfer system. Following transfer, membranes were blocked with 5% milk in TBST before incubating with a designated primary antibody (all diluted 1:1000 in 2% BSA in TBST, except PFKFB3 at 1:2000 in 2% BSA in TBST) overnight at 4°C. All antibodies utilized in this study are listed in Table S5. After primary incubation, membranes were washed 3 times with TBST buffer and then incubated with anti-rabbit IgG, HRP-linked secondary antibody (diluted 1:10,000 in 5% milk in TBST; Cell Signaling Technologies #7074) for 1 hour at 25°C. After secondary incubation, membranes were washed 3 times with TBST before detection. Membranes were developed using SuperSignal™ West Pico PLUS Chemiluminescent Substrate (Thermo Fisher Scientific #34577) and imaged with iBright 1500 (Thermo Fisher Scientific).

### Cellular F26BP Extraction

2 x 10^6^ cells were seeded per 10 cm dish in 4 mL of cDMEM. Twenty-four hours after seeding, 4 mL of media was added per well supplemented with doxycycline (final: 300 ng/mL; Sigma #D9891) or water. Twenty-four hours later, media was replaced with 7 mL of minimal DMEM (US Biological Life Sciences #D9800-28, see Table S4 for media composition) supplemented with 5 mM glucose, 4 mM glutamine, 3.7 g/L NaHCO_3_ (pH 7.4), and 10% dialyzed FBS (Thermo Fisher Scientific #26400044) with 1 μM Antimycin A (Millipore Sigma #A8674) or dimethylsulfoxide (DMSO; Thermo Fisher Scientific #D12345). After 1 hour incubation in new media, F26BP was extracted, removing media, adding 250 μL 0.1 M NaOH, and cooking extracts at 90°C for 10 minutes. To normalize measurements, cellular protein in F26BP extracts was quantified by Pierce™ BCA protein assay kit (Thermo Fisher Scientific #23227).

### Measurement of Oxygen Consumption Rate

20-30 x 10^3^ cells were seeded per well of XFe24 24-well microplates in 200 μL of cDMEM. Four hours after seeding, an additional 300 μL of cDMEM was added to each well (total volume per well: 500 μL). Twenty-four hours later, 500 μL of media was added per well supplemented with doxycycline (final: 300 ng/mL; Sigma #D9891) or water. Twenty-four hours later, media was replaced with 500 μL of minimal DMEM (US Biological Life Sciences #D9800-28, see Table S4 for media composition) supplemented with 5 mM glucose, 4 mM glutamine, 5 mM HEPES (pH 7.4), and 10% dialyzed FBS (Thermo Fisher Scientific #26400044). Extracellular acidification rates (ECAR) and oxygen consumption rates (OCR) were measured using Agilent Seahorse XFe24 Analyzer. Four basal measurements were performed for 4 minutes after a 2-minute mixing and 30-second waiting period. Four maximal glycolytic measurements were collected upon the addition of 1 μM Antimycin A (Millipore Sigma #A8674) for 4 minutes a 2-minute mixing and 2-minute waiting period. To normalize measurements by cellular protein, cells were lysed with 50 μL of a lysis buffer (50 mM Tris (pH 8), 1 mM EDTA, 1% SDS) and protein content was measured by Pierce™ BCA protein assay kit (Thermo Fisher Scientific #23227).

### Measurement of Lactate Secretion Rate

0.2 x 10^6^ cells were seeded per well of 6-well plate in 2 mL of cDMEM. Twenty-four hours after seeding, 2 mL of cDMEM was added per well supplemented with doxycycline (final: 300 ng/mL; Sigma #D9891) or water. Twenty-four hours later, media was replaced with minimal DMEM (US Biological Life Sciences #D9800-28, see Table S4 for media composition) supplemented with 5 mM glucose, 4 mM glutamine, 3.7 g/L NaHCO_3_ (pH 7.4), and 10% dialyzed FBS (Thermo Fisher Scientific #26400044) – allowing cells to adjust to new media for 1 hour. After incubation, media was exchanged to 2 mL of the experimental media with either 1 μM Antimycin A (Millipore Sigma #A8674) or DMSO (Thermo Fisher Scientific #D12345). Thereafter, 150 μL spent media samples were collected at 2, 4, and 8 hours and immediately frozen on dry ice. Samples were stored at - 80°C until analysis. After measurements, cells were lysed with 150 μL of a lysis buffer (50 mM Tris (pH 8), 1 mM EDTA, 1% SDS) and protein content was measured by Pierce™ BCA protein assay kit (Thermo Fisher Scientific #23227). Lactate was quantified using Colorimetric L-Lactate Assay Kit-I (Eton Bioscience #120001400A) per the manufacturer’s instructions. Linear regression was performed across the 8-hour time course to determine the lactate secretion and glucose uptake rates.

### Estimation of ATP Production Rate from Experimental Glycolysis and Respiration Rates

ATP production rates for glycolysis and respiration were calculated using lactate secretion rate (LSR) and oxygen consumption rate (OCR) using the ATP yields from a previous study^60^. Briefly, it is widely accepted that fermentative glycolysis yields two molecules of ATP and two molecules of lactate per molecule of glucose. Therefore, for glycolytic ATP production rates were equated to the measured lactate secretion rates.

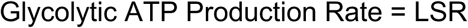

For mammalian respiration, the estimates needed to account for the ATP directly produced from both glycolysis and the TCA cycle, H^+^ pump efficiency of the electron transport chain (ETC), and the mitochondrial proton leak. Experimentally, the respiration ATP production rates derived from the measured oxygen consumption rates (OCR)^61^. To account for the mitochondrial proton leak, OCR in the presence of antimycin A was first subtracted from basal OCR. Then, based on the number of ATP and reducing equivalents directly produced, number of protons pumped per electron, and the number of ATP yielded per proton, it was assumed 24 molecules of ATP were generated per molecule of oxygen consumed. To summarize this conversion:

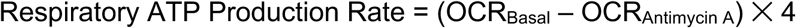

### Expression and Purification of PFKFB3

cDNA encoding for human PFKFB3 (NCBI GenBank mRNA transcript variant 1, Reference Sequence: NM_004566.4) was subcloned into pET-28a(+) (Novagen #69864) within the multiple cloning region at the *NdeI* and *EcoRI* restriction sites. The N and C-terminal 6His-tags were modified using Q5 Site-Directed Mutagenesis Kit (New England Biolabs #E0554). The N-terminal 6His-tag was modified to: (1) insert a PreScission (HRV-3C) cleavage site (Pre) and (2) account for eukaryotic co-translational methionine cleavage. These N-terminal alterations were designed such that PreScission (underlined) would cleave (/) just before the endogenous PFKFB3 N-terminus (bold) (HHHHHHSSGLVPRGGGSGGGS<u>LEVLFQ/G**P**</u>**LELTQSRVQKI**…). The C-terminal 6His-tag was fully removed such that the PFKFB3 stop codon was immediately upstream of the T7 terminator sequence. Construct accuracy was verified using DNA sequencing.

pET28a-His-Pre-PFKFB3 transformed Rosetta 2 (DE3) pLysS competent cells (Millipore Sigma #71401) were grown in Fisher BioReagents™ Terrific Broth (+ 0.4% glycerol; Fisher Scientific #BP2468500) at 37°C until an OD_600nm_ of 0.6-0.8. At this point, recombinant protein expression was induced by treating with 0.4 mM ITPG (Fisher Scientific #BP1620) and incubating at 16°C for 16 hours. Unless otherwise specified, all recombinant protein purification steps took place at 4C. To lyse, the bacterial pellet was resuspended in lysis buffer (final: 50 mM Tris (pH 8.0), 0.3 M NaCl, 15 mM imidazole, 10 mM Na_2_HPO_4_, 1 mM TCEP, 1 mg/mL lysozyme, 20 μg/mL DNaseI, 5 mM MgSO_4_, and Pierce Protease Inhibitor Tablet (Thermo Fisher Scientific #A32963)) and treated with B-Per™ II Bacterial Protein Extraction Reagent (Thermo Fisher Scientific #78260) in a 1:1 ratio (final: 2 g pellet/mL total lysis volume) for 20 minutes at 25°C.

Cleared lysate was applied to HisPur™ Ni-NTA Resin (Thermo Fisher Scientific #88221) and eluted with an elution buffer (50 mM Tris (pH 8.0), 0.3 M NaCl, 300 mM imidazole, 10 mM Na_2_HPO_4_, 1 mM TCEP). The eluant underwent a quick buffer exchange to buffer A (50 mM Tris (pH 8), 20 mM KCl, 5 mM KH_2_PO_4_, 1 mM TCEP, 0.1 mM EDTA, and 10% glycerol) via Zeba™ Spin Desalting Columns (40K MWCO; Thermo Fisher Scientific #A4002471). The concentrate was treated with PreScission (roughly 1 U/100 μg PFKFB3; GenScript #Z03092) at 4°C for 1 hour. Cleaved eluant was applied to HiTrap™ Q HP (1 mL; Cytiva #17115301) and eluted with a 20-500 mM KCl linear gradient via the ÄKTA start™ chromatography system equipped with Frac30 fraction collector and UNICORN™ start control software. Eluant fractions containing PFKFB3 were dialyzed against storage buffer A (50 mM Tris (pH 8), 50 mM KCl, 1 mM TCEP, 0.1 mM EDTA, 20% glycerol) using Slide-A-Lyzer™ G3 Dialysis Cassettes (20K MWCO; Thermo Fisher Scientific #A52976), and stored at −80°C. Protein concentrations were quantified by Pierce™ Bradford Plus Protein Assay Reagent (Thermo Fisher Scientific #23238), and purity was evaluated by SDS-PAGE via Imperial™ Protein Stain (Thermo Fisher Scientific #24615).

### PFKFB Activity

In a final reaction volume of 100 μL, 100 μg/mL PFKFB3 was incubated with 0.2 mM F6P, 3 mM ATP coordinated with equimolar MgCl_2_, and effectors as indicated for 20 minutes at 37°C in reaction buffer A (50 mM HEPES (pH 7.4), 100 mM KCl, 1 mg/ml BSA, 0.1 mM EDTA, 5 mM MgCl_2_, and 1 mM DTT). No F6P was included as a negative control. For metabolite screen, all regulators had a final concentration of 1 mM, except Acetyl-CoA, Malonyl-CoA, Oleoyl-CoA and Palmityl-CoA at 0.1 mM. For inhibitor titration, regulators were added at the indicated concentrations. All metabolites included in this study are listed in Table S3. Reactions were quenched by adding NaOH (final: 0.2 M) and heating the mixture to 90°C for 10 minutes. Samples were stored at −20°C until analysis. In addition to F26BP quantification, PFKFB3 activity was measured using Fructose-6-Phosphate Fluorometric Assay Kit (Abcam #ab204720) and ADP-Glo™ Kinase Assay Kit (Promega #V6930) per the manufacturer’s instructions.

### F26BP Quantification

F26BP in cell extracts and *in vitro* PFKFB3 activity assays were measured using the potato pyrophosphate-dependent 6-phosphofructokinase (PP_i_ PFK) coupled enzyme assay previously described^62^. All purification steps and buffers were maintained at 4°C. PP_i_ PFK was purified from peeled and washed *Potato Tubers* by blending in a 1:1 wt/vol with lysis buffer (20 mM HEPES (pH 8.2), 20 mM potassium acetate, 1 mM EDTA, 2 mM TCEP, and Pierce Protease Inhibitor Tablets (Thermo Fisher Scientific #A32963)). Blended solution was cleared by passing through 4 layers of cheese cloth, and centrifuging the filtrate at 3,000g for 10 minutes. The supernatant was collected and treated with 80% ammonium sulfate in a 1:1 vol/vol (final: 40% ammonium sulfate) to precipitate PP_i_ PFK. PP_i_ PFK was isolated by centrifuging 3,000g for 30 minutes and resuspending the resultant pellet in storage buffer B (20 mM Tris (pH 8.2), 20 mM KCl, 2 mM TCEP, and 20% glycerol; 1/10 precipitation volume). The resuspension underwent dialysis against storage buffer B using Slide-A-Lyzer™ G3 Dialysis Cassettes (10K MWCO; Thermo Fisher Scientific #A52972), and stored at 4°C for up to two weeks.

F26BP was measured by adapting the PP_i_ PFK coupled enzyme assay to a 96 well plate format (200 μL per well). Briefly, each reaction comprised of 20 μL sample dilution, 40 μL PP_i_ PFK, 0.1 U Aldolase (Sigma Aldrich #A8811), 0.5 U Triose Phosphate Isomerase (Millipore Sigma #T2391), and 0.5 U Glycerol-3-Phosphate Dehydrogenase (Roche #10127752001), 5 mM F6P, 0.15 mM NADH, and 2.5 mM pyrophosphate in reaction buffer B (50 mM Tris (pH 8), 5 mM MgCl_2_, and 1 mM DTT). Commercial auxiliary enzymes were desalted with a 30K MWCO centrifugal filter (Millipore #UFC8030) in reaction buffer B prior to use. The reaction was initiated with the addition of pyrophosphate. Immediately following initiation, NADH consumption was monitored spectrophotometrically at 340 nm for 2 minutes (measurement frequency: 6 seconds^-1^) at 30°C via BioTek Cytation1. Reaction rates were first quantified by taking the slope of the longest linear region (minimum: 1 minute), and then divided by the rate of the No F26BP control sample. Due to the lack of a F26BP standard, all samples underwent a logarithmic serial dilution in their respective buffers to construct a EC_50_ curve for PP_i_ PFK. The top and bottom of EC_50_ curves were constrained for all samples, and defined by positive and negative sample controls. The sample dilution that achieved PP_i_ PFK EC_50_ was equated to 5 nM F26BP and utilized to calculate F26BP content in the original sample^62^.

#### In Vitro Glycolysis

Human PFKP expression, purification, and enzyme activity assays were performed as previously described^47,63^. Briefly, the F26BP regulatory circuit was continuously monitored by using a partial reconstitution of upper glycolysis enzymes in a 96-well plate (200 μL per well). Each reaction contained 0.13 U Aldolase (Millipore Sigma #A8811), 1 U Triose Phosphate Isomerase (Millipore Sigma #T2391), 0.4 U Glycerol-3-Phosphate Dehydrogenase (Roche #10127752001), 0.2 mM F6P, 3 mM ATP chelated with equimolar MgCl_2_, 0.3 mM ADP chelated with equimolar MgCl_2_, 1 mM Phosphate, and 0.15 mM NADH in reaction buffer A (50 mM HEPES (pH 7.4), 100 mM KCl, 1 mg/ml BSA, 0.1 mM EDTA, 5 mM MgCl_2_, and 1 mM DTT). Regulators were added at the indicated concentrations. Commercial auxiliary enzymes were desalted with a 30K MWCO centrifugal filter (Millipore #UFC8030) in reaction buffer A prior to use. To initiate F26BP production, 10 μg/mL of PFKFB3 was pre-incubated in reaction mixture for 20 minutes at 37°C. After pre-incubation, 1.61 μg/mL purified human PFKP was added to the reaction mixture to initiate *in vitro* glycolysis reaction. Immediately following initiation, NADH consumption was monitored spectrophotometrically at 340 nm for 60 minutes (measurement frequency: 13 seconds^-1^) at 37°C via BioTek Cytation1. Reaction rates were first quantified by taking the slope of the longest linear region (minimum: 10 minutes), and then divided by the rate of the No F6P control sample.

## CONTRIBUTIONS

Conceptualization, M.M.K. and D.V.T; methodology, M.M.K., X.Y. and D.V.T.; human phosphofructokinase purification B.A.W.; data curation, M.M.K., X.Y., and D.V.T.; funding acquisition and supervision, D.V.T.; writing - original draft, M.M.K. and D.V.T; writing - review & editing, M.M.K., B.A.W., and D.V.T.

## ACKNOWLEDGEMENTS

We graciously thank Yong-Hwan Lee for supporting method development and providing feedback throughout the project, as well as members of the Titov lab for many helpful suggestions along the way. Research reported in this publication was supported by the National Institute of General Medical Sciences of the National Institutes of Health under award number R35 GM152114 to D.V.T., the Visual Sciences CoBRE project leader funding (P20GM144230) to B.A.W., and National Institute of General Medical Sciences funding (R35GM158392) to B.A.W.

## SUPPLEMENTAL TABLES

**Table S1.**
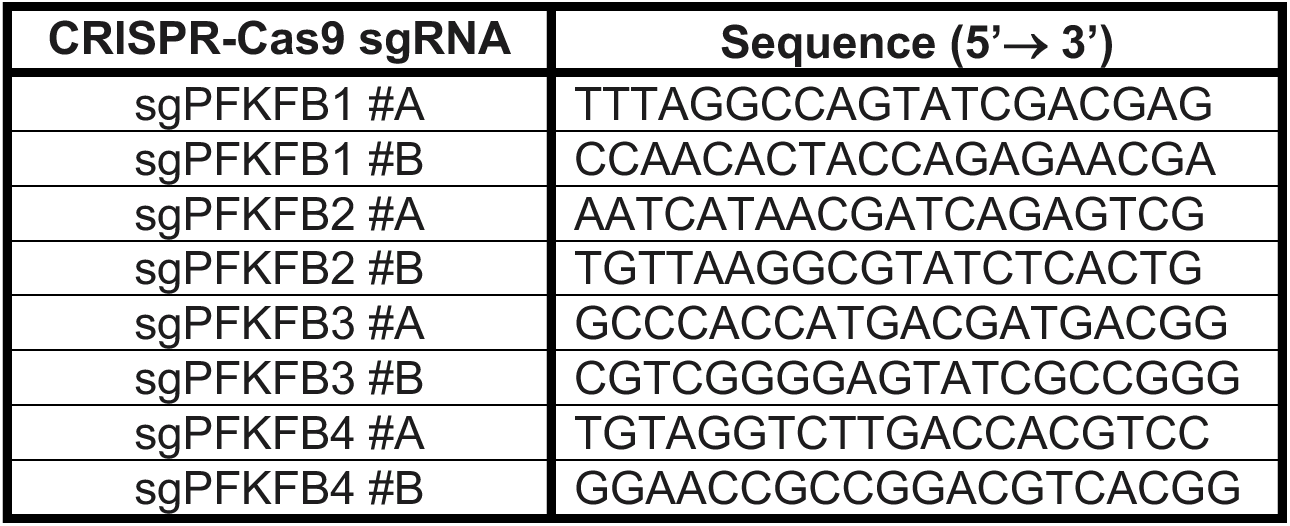
Guide RNA Sequences used to generate *PFKFB1-4* knockout (PFKFB_0_)

**Table S2.**
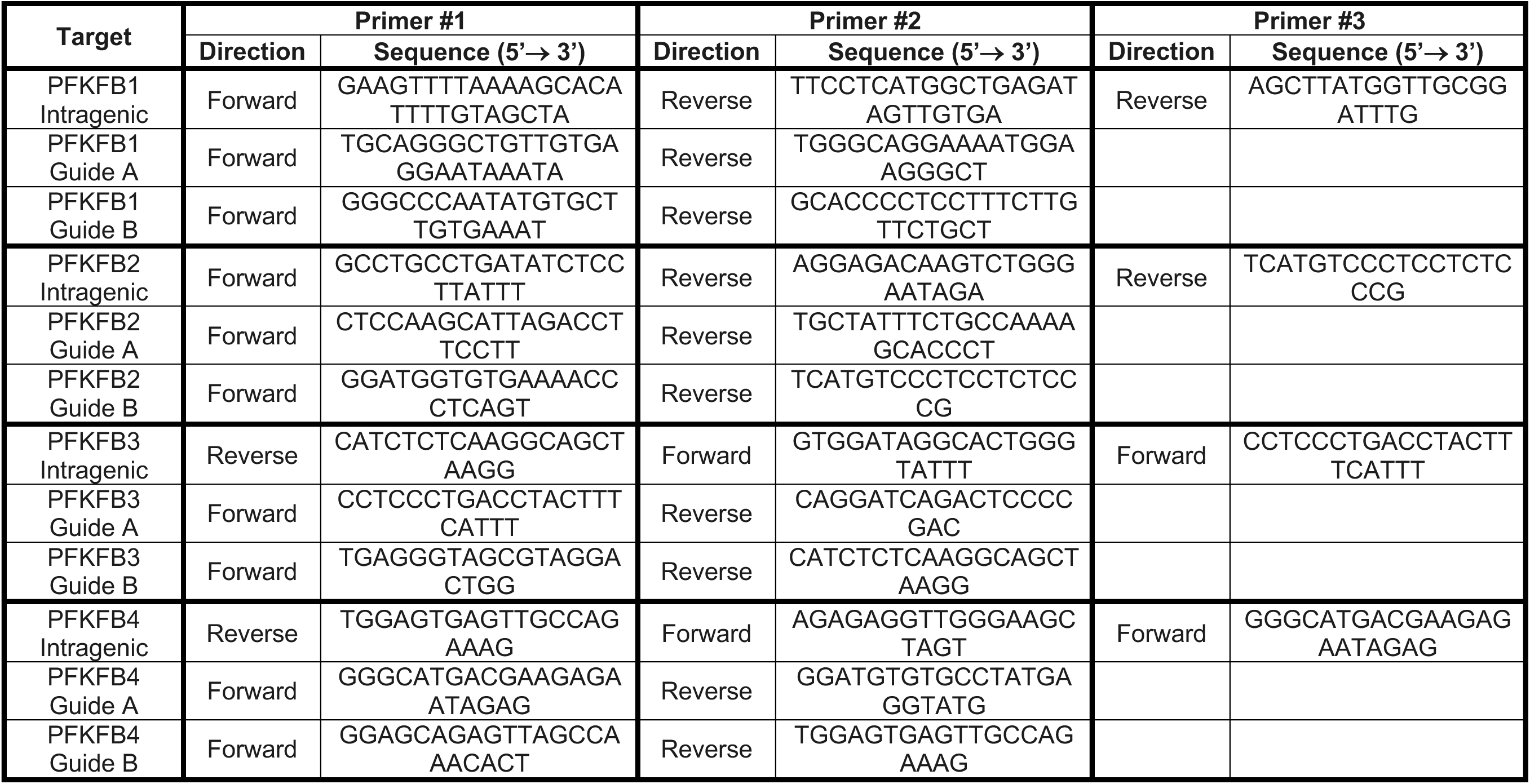
PCR and Sequencing Primers used to evaluate *PFKFB1-4* Insertion/Deletion & Intragenic Deletion Frequency.

**Table S3.** Metabolites utilized in PFKFB3 Regulator Screen.

| Chemical | Source | Cat# |
| --- | --- | --- |
| Adenosine 5'-triphosphate disodium salt hydrate (ATP) | Millipore Sigma | A6419 |
| Potassium Phosphate Monobasic (KH<sub>2</sub>PO<sub>4</sub>) | Fisher Scientific | P285 |
| Adenosine 5'-diphosphate sodium salt (ADP) | Millipore Sigma | A2754 |
| Adenosine 5'-monophosphate monohydrate (AMP) | Millipore Sigma | A2252 |
| D-Glucose 6-phosphate sodium salt (G6P) | Millipore Sigma | G7879 |
| α-D-Glucose 1-phosphate disodium salt hydrate (G1P) | Millipore Sigma | G7000 |
| D-Fructose-6-phosphate disodium salt (F6P) | Thermo Scientific | J66311.03 |
| D-Fructose 1,6-bisphosphate trisodium salt hydrate (F16BP) | Millipore Sigma | F6803 |
| Dihydroxyacetone phosphate dilithium salt (DHAP) | Millipore Sigma | D7137 |
| DL-Glyceraldehyde 3-phosphate solution (GAP) | Millipore Sigma | G5251 |
| D-(-)-3-Phosphoglyceric acid disodium salt (3PG) | Millipore Sigma | P8877 |
| L-2-Phosphoglyceric acid disodium salt hydrate (2PG) | Millipore Sigma | 19710 |
| Phosphoenol-pyruvate (PEP) | Millipore Sigma | 10108294001 |
| Sodium Pyruvate | Thermo Scientific | 11360070 |
| Sodium L-Lactate | Millipore Sigma | 71718 |
| α-Ketoglutaric acid (α-KG) | Millipore Sigma | 75890 |
| D-α-Hydroxyglutaric acid sodium salt | Cayman Chemical | C832G02 |
| Succinic acid | Thermo Scientific | 219552500 |
| Fumaric acid | Millipore Sigma | 47910 |
| Itaconic acid | Millipore Sigma | I29204 |
| cis-Aconitic acid | Millipore Sigma | A3412 |
| Sodium citrate tribasic dihydrate | Millipore Sigma | C8532 |
| L-(-)-Malic acid | Millipore Sigma | 02288 |
| Acetyl coenzyme A lithium salt (Acetyl CoA) | Millipore Sigma | A2181 |
| Oxaloacetic acid (OAA) | Millipore Sigma | O4126 |
| Malonyl coenzyme A lithium salt (Malonyl CoA) | Millipore Sigma | M4263 |
| Oleoyl coenzyme A lithium salt (Oleoyl CoA) | Millipore Sigma | O1012 |
| Palmitoyl coenzyme A lithium salt (Palmitoyl CoA) | Millipore Sigma | P9716 |
| sn-Glycerol 3-phosphate lithium salt (G3P) | Millipore Sigma | P94124 |
| L-Serine | Millipore Sigma | S4500 |
| L-Alanine | Millipore Sigma | A7627 |
| L-Glycine | Millipore Sigma | 50046 |
| L-Glutamine | Millipore Sigma | G8540 |
| L-Leucine | Millipore Sigma | L8000 |
| 6-Phosphogluconic Acid Trisodium Salt Dihydrate | Millipore Sigma | P7877 |
| D-Ribulose 5-phosphate sodium salt (Ribulose-5P) | Millipore Sigma | R9875 |
| D-Ribose 5-phosphate disodium salt hydrate (Ribose-5P) | Millipore Sigma | R7750 |
| D-Xylulose 5-phosphate lithium salt (Xylulose-5P) | Millipore Sigma | 15732 |
| D-Sedoheptulose 7-phosphate lithium salt (Sedoheptulose-7P) | Millipore Sigma | 78832 |
| D-Erythrose 4-phosphate sodium salt (Erythrose-4P) | Millipore Sigma | E0377 |
| β-Nicotinamide adenine dinucleotide phosphate hydrate (NADP) | Millipore Sigma | N5755 |
| β-Nicotinamide adenine dinucleotide 2'-phosphate reduced tetrasodium salt hydrate (NADPH) | Millipore Sigma | N6505 |
| β-Nicotinamide adenine dinucleotide hydrate (NAD) | Millipore Sigma | N1636 |
| β-Nicotinamide adenine dinucleotide, reduced disodium salt hydrate (NADH) | Millipore Sigma | N8129 |

**Table S4.** Cell Culture Media Composition.

| **Component (mg/L)** | **Standard DMEM**<br>**(Gibco #11995)** | **Minimal DMEM**<br>**(US Biological Life**<br>**Sciences #D9800-D28)** |
| --- | --- | --- |
| Glycine | 30 | 0 |
| L-Arginine hydrochloride | 84 | 0 |
| L-Cystine 2HCl | 63 | 0 |
| L-Glutamine | 584 | 0 |
| L-Histidine hydrochloride-H<sub>2</sub>O | 42 | 0 |
| L-Isoleucine | 105 | 0 |
| L-Leucine | 105 | 0 |
| L-Lysine hydrochloride | 146 | 0 |
| L-Methionine | 30 | 0 |
| L-Phenylalanine | 66 | 0 |
| L-Serine | 42 | 0 |
| L-Threonine | 95 | 0 |
| L-Tryptophan | 16 | 0 |
| L-Tyrosine disodium salt dihydrate | 104 | 0 |
| L-Valine | 94 | 0 |
| Choline chloride | 4 | 4 |
| D-Calcium pantothenate | 4 | 4 |
| Folic Acid | 4 | 4 |
| Niacinamide | 4 | 4 |
| Pyridoxine hydrochloride | 4 | 4 |
| Riboflavin | 0.4 | 0.4 |
| Thiamine hydrochloride | 4 | 4 |
| i-Inositol | 7.2 | 7.2 |
| Calcium Chloride (CaCl<sub>2</sub>) (anhyd.) | 200 | 265 |
| Ferric Nitrate (Fe(NO<sub>3</sub>)<sub>3</sub>·9H<sub>2</sub>O) | 0.1 | 0.1 |
| Magnesium Sulfate (MgSO<sub>4</sub>) (anhyd.) | 97.67 | 97.67 |
| Potassium Chloride (KCl) | 400 | 400 |
| Sodium Bicarbonate (NaHCO<sub>3</sub>) | 3700 | 0 |
| Sodium Chloride (NaCl) | 6400 | 6400 |
| Sodium Phosphate monobasic<br>(NaH<sub>2</sub>PO<sub>4</sub>·H<sub>2</sub>O) | 125 | 109 |
| D-Glucose (Dextrose) | 4500 | 0 |
| Phenol Red | 15 | 15.9 |
| Sodium Pyruvate | 110 | 0 |

**Table S5.** Primary & Secondary Antibodies used for Western Blotting.

| **Antibody** | **Source** | **Identifier** |
| --- | --- | --- |
| Rabbit polyclonal 6-phosphofructo-2-kinase/fructose-2,6-biphosphatase 3 (PFKFB3) | Proteintech | Cat#13763-1-AP; RRID: AB_2162854 |
| Rabbit monoclonal 6-phosphofructo-2-kinase/fructose-2,6-biphosphatase 2 (PFKFB2) | Cell Signaling Technologies | Cat#13045; RRID: AB_2798097 |
| Rabbit polyclonal DYKDDDDK (Flag) Tag | Cell Signaling Technologies | Cat#2368; RRID: AB_2217020 |
| Rabbit polyclonal Firefly Luciferase | Thermo Fisher Scientific | Cat#PA5-32209; RRID: AB_2549682 |
| Rabbit monoclonal β-Actin | Cell Signaling Technologies | Cat#4970; RRID: AB_2223172 |
| Rabbit monoclonal Hexokinase 1 (HK1) | Cell Signaling Technologies | Cat#2024; RRID: AB_2116996 |
| Rabbit monoclonal Hexokinase 2 (HK2) | Cell Signaling Technologies | Cat#2867; RRID: AB_2232946 |
| Rabbit monoclonal Phosphofructokinase platelet isozyme (PFKP) | Cell Signaling Technologies | Cat#8164; RRID: AB_2713957 |
| Rabbit monoclonal Glyceraldehyde-3-phosphate dehydrogenase (GAPDH) | Cell Signaling Technologies | Cat#2118; RRID: AB_561053 |
| Rabbit monoclonal Pyruvate kinase M2 isoform (PKM2) | Cell Signaling Technologies | Cat#4053; RRID: AB_1904096 |
| Rabbit monoclonal Mitochondrial pyruvate carrier 1 (MPC1) | Cell Signaling Technologies | Cat#14462; RRID: AB_2773729 |
| Rabbit monoclonal Mitochondrial pyruvate carrier 2 (MPC2) | Cell Signaling Technologies | Cat#46141; RRID: AB_2799295 |
| Rabbit monoclonal Citrate Synthase | Cell Signaling Technologies | Cat#14309; RRID: AB_2665545 |
| Rabbit monoclonal Aconitase 2 (ACO2) | Cell Signaling Technologies | Cat#6571; RRID: AB_2797630 |
| Rabbit polyclonal Isocitrate dehydrogenase 1 (IDH1) | Cell Signaling Technologies | Cat#3997 |
| Rabbit monoclonal Isocitrate dehydrogenase 2 (IDH2) | Cell Signaling Technologies | Cat#56493; RRID: AB_2799511 |
| Goat anti-rabbit IgG, HRP-linked antibody | Cell Signaling Technologies | Cat#7074; RRID: AB_2099233 |

## SUPPLEMENTAL DATA

**Figure S1.**
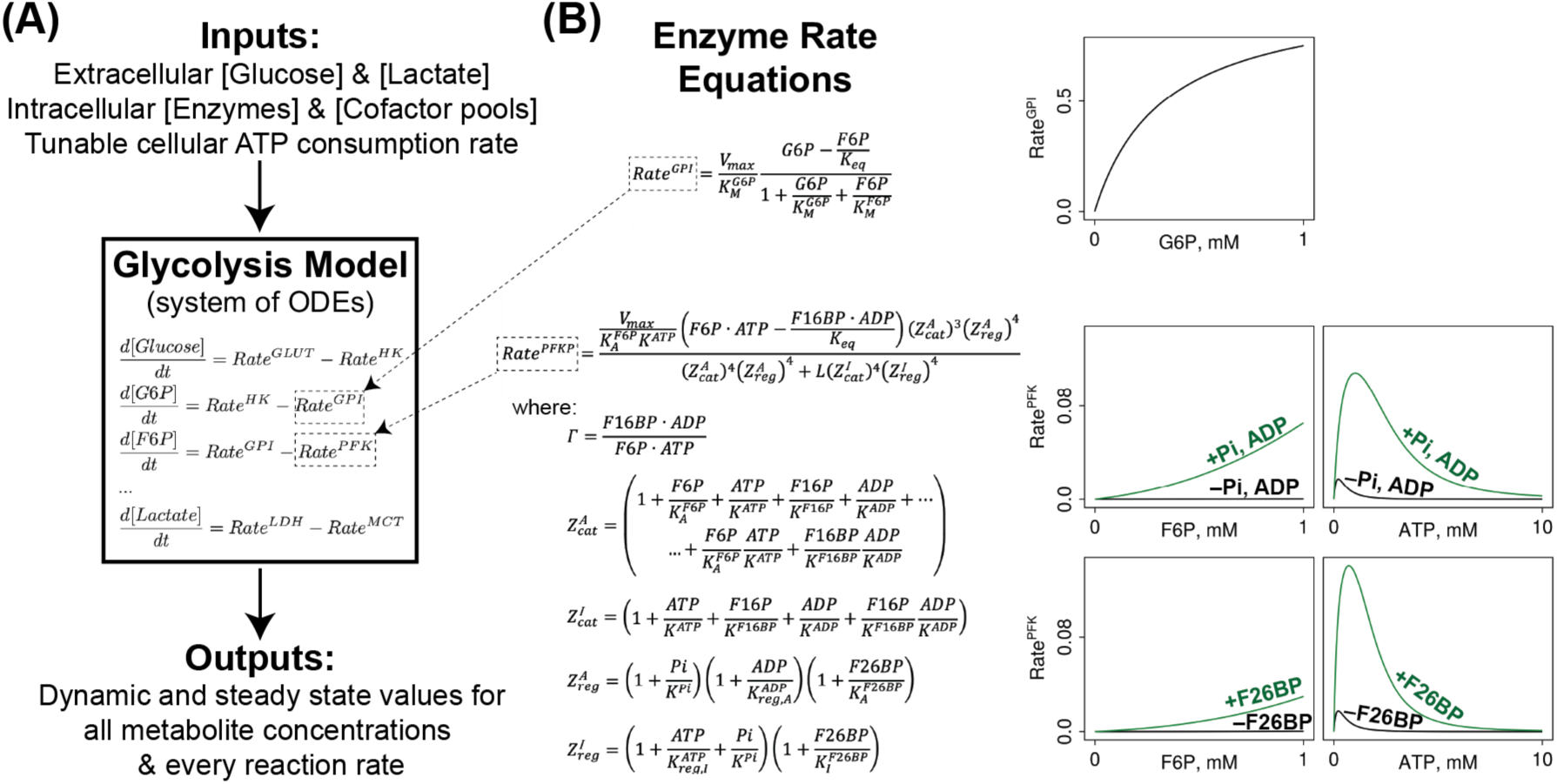
Biophysical model of mammalian glycolysis overview. (A) Schematic of the glycolysis inputs and outputs. (B) Kinetic rate equations are shown for GPI and PFK, with special attention to allosteric regulation by F26BP. Plots are actual GPI and PFK rates calculated by the respective equations with rates normalized by the V_max_. Note the dramatic allosteric activation of the PFK rate in the presence of F26BP.^1^

**Figure S2.**
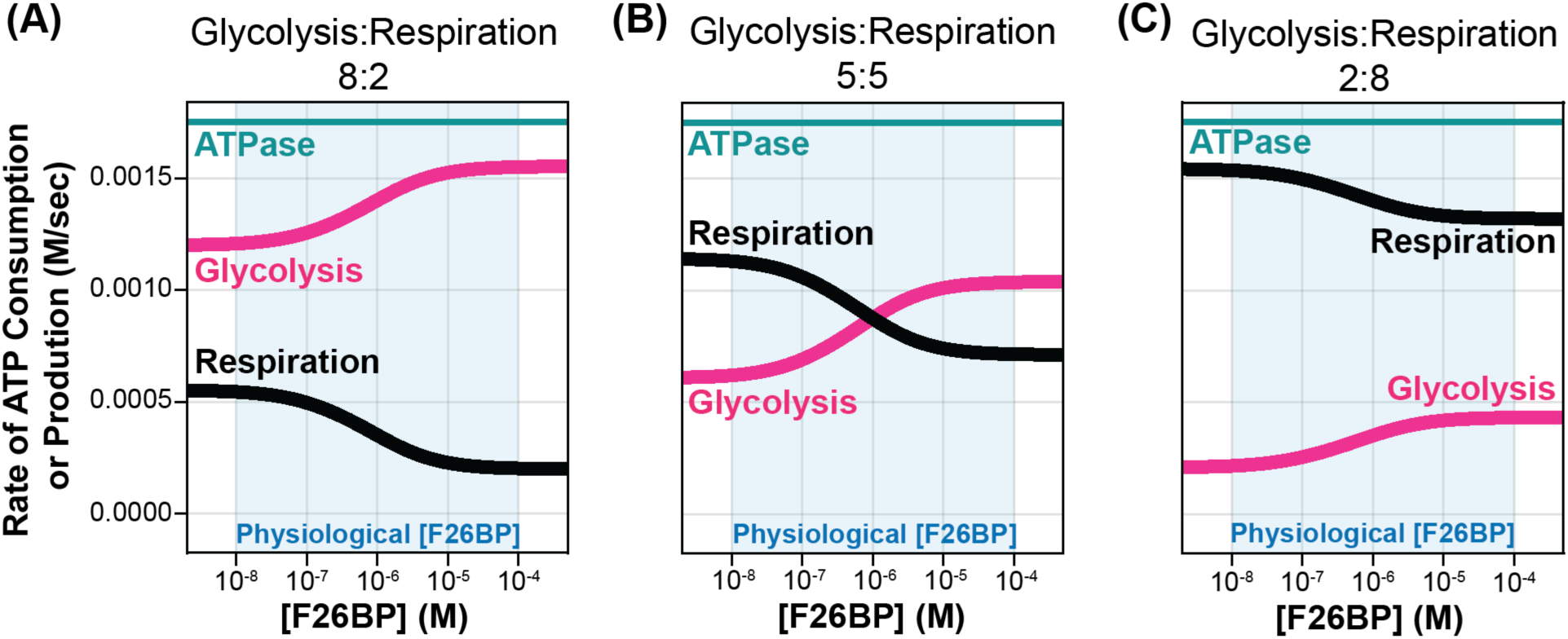
Effect of F26BP on glycolytic and respiratory ATP production rates at varying respiration-to-glycolysis ratios. Steady-state simulations of a glycolysis model with added ATP Synthase to mimic mitochondrial respiration were performed across a range of F26BP concentrations (2×10^-9^ to 5×10^-4^ M) while varying the ratio of *V_max_* of glycolysis and respiration: (A) 8:2, (B) 5:5, and (C) 2:8. The glycolytic ATP production rate is represented by PKM2 flux (pink), respiration rate by ATP synthase flux (black), and ATP consumption rate by ATPase flux (teal). The sum of *V_max_* of glycolysis and respiration was kept constant. ATP consumption (ATPase Vmax) was set to 20% of total ATP supply capacity and held constant across all conditions. The light blue shaded region indicates the physiological F26BP concentration range (10^-8^ to 10^-4^ M).

**Figure S3.**
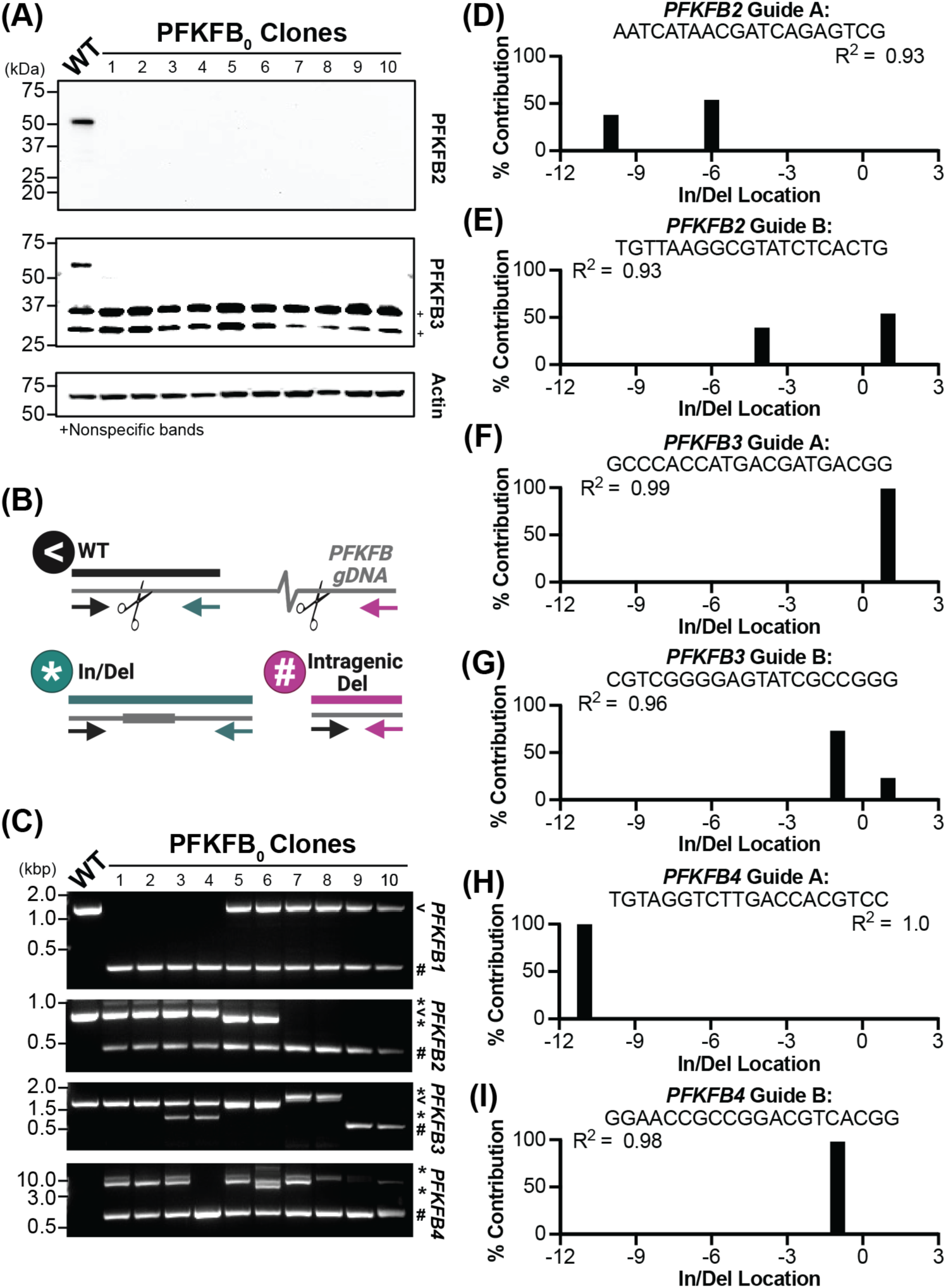
Generation and validation of clonal PFKFB_0_ knock outs. (A) Western blot showing endogenous PFKFB2 and PFKFB3 expression in HeLa (wildtype, WT) and 10 genetically independent PFKFB_0_ clones. PFKFB_0_ clone 1 was utilized for further cell line derivation and investigations throughout this study. Representative gel from one of two independent experiments. (B) Schematic of CRISPR/Cas9 gRNA (scissor) and primer design (arrows), which generated and evaluated insertion/deletion (in/del) and intragenic deletions in PFKFB genes, respectively. (C) PCR amplification of *PFKFB1, PFKFB2, PFKFB3,* and *PFKFB4* genes within WT and 10 genetically distinct PFKFB_0_ clones. Resolved PCR products are labeled in accordance with B schematic: WT (<), local in/del at amplified cut site (*), and severe intragenic deletion (#). (D-I) Monoallelic knock out efficiency of PFKFB_0_ clone 1 as evaluated by EditCo’s Inference of CRISPR Edits (ICE)^58,59^. Estimates the contribution of insertion/deletions at gRNA targeting: (D) site A within *PFKFB2,* (E) site B within *PFKFB2,* (F) site A within *PFKFB3,* (G) site B within *PFKFB3,* (H) site A within *PFKFB4,* and (I) site B within *PFKFB4.* R^2^ denotes Pearson correlation coefficient, indicating how well the ICE model fits with the Sanger Sequencing data.

**Figure S4.**
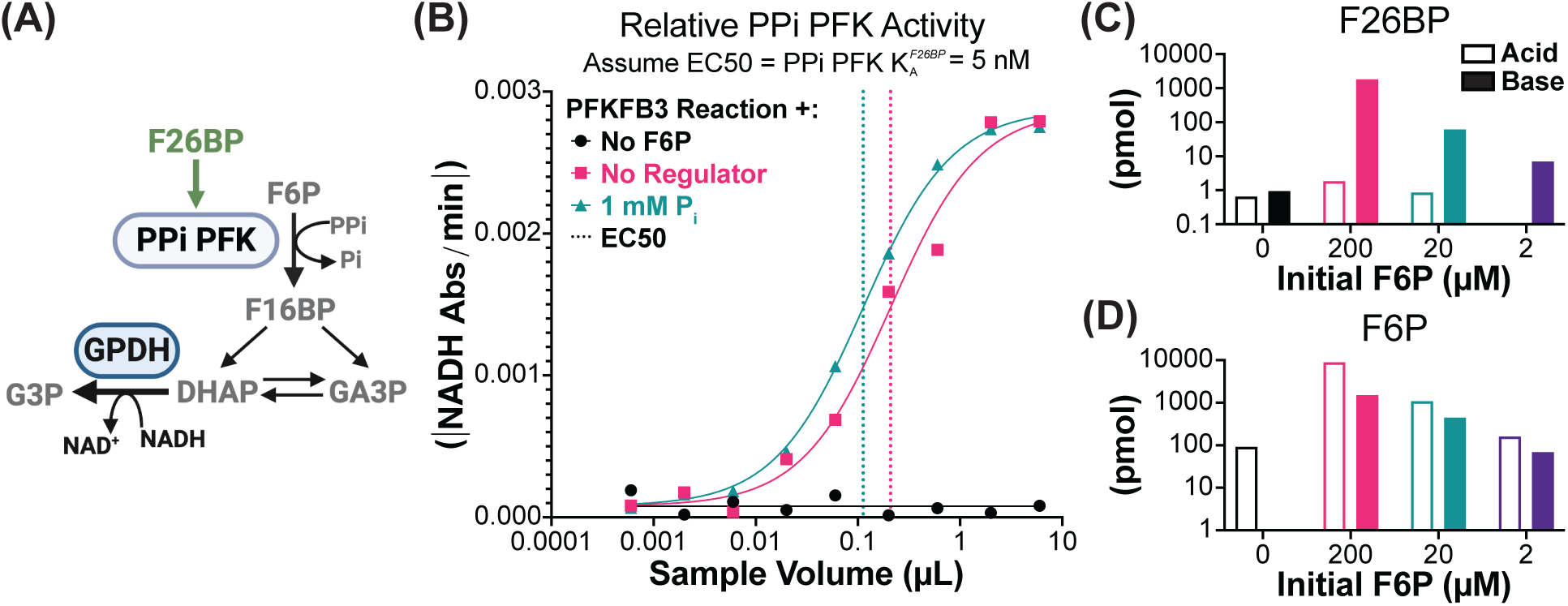
F26BP quantification method. (A) Schematic of the potato pyrophosphate-dependent 6-phosphofructokinase (PP_i_ PFK) coupled enzyme assay previously described^62^. Notable intermediates (grey text), enzymes (grey ovals), and allosteric regulators (green text) are highlighted. Coupled enzyme reaction terminates with glycerol-3-phosphate dehydrogenase (GPDH), which allows for continuous monitoring by NADH consumption (black text). (B) Representative PP_i_ PFK EC_50_ curve (solid line) from PFKFB3 reaction serial dilution series. PFKFB3 reaction conditions included: no F6P (black, negative control), no regulator (pink), and the addition of 1 mM P_i_ (teal, positive control). PP_i_ PFK EC_50_ (dashed line) was equated to 5 nM F26BP and utilized to calculate F26BP in sample^62^. (C-D) Validating F26BP quantification by quenching PFKFB3 reaction with either acid (outline; F26BP-destabilizing negative control) or base (solid; F26BP-stabilizing positive control). PFKFB3 reaction was incubated with 3 mM ATP and 0 μM (black), 200 μM (pink), 20 μM (teal), or 2 μM (purple) F6P. Measurement from one independent experiment. (C) F26BP content estimated via PP_i_ PFK activity assay. (D) F6P content quantified via commercial F6P fluorometric kit. In cross comparing F6P utilized to F26BP produced, differences between the acid and base quenching methods correlated with the amount of F26BP stabilized.

**Figure S5.**
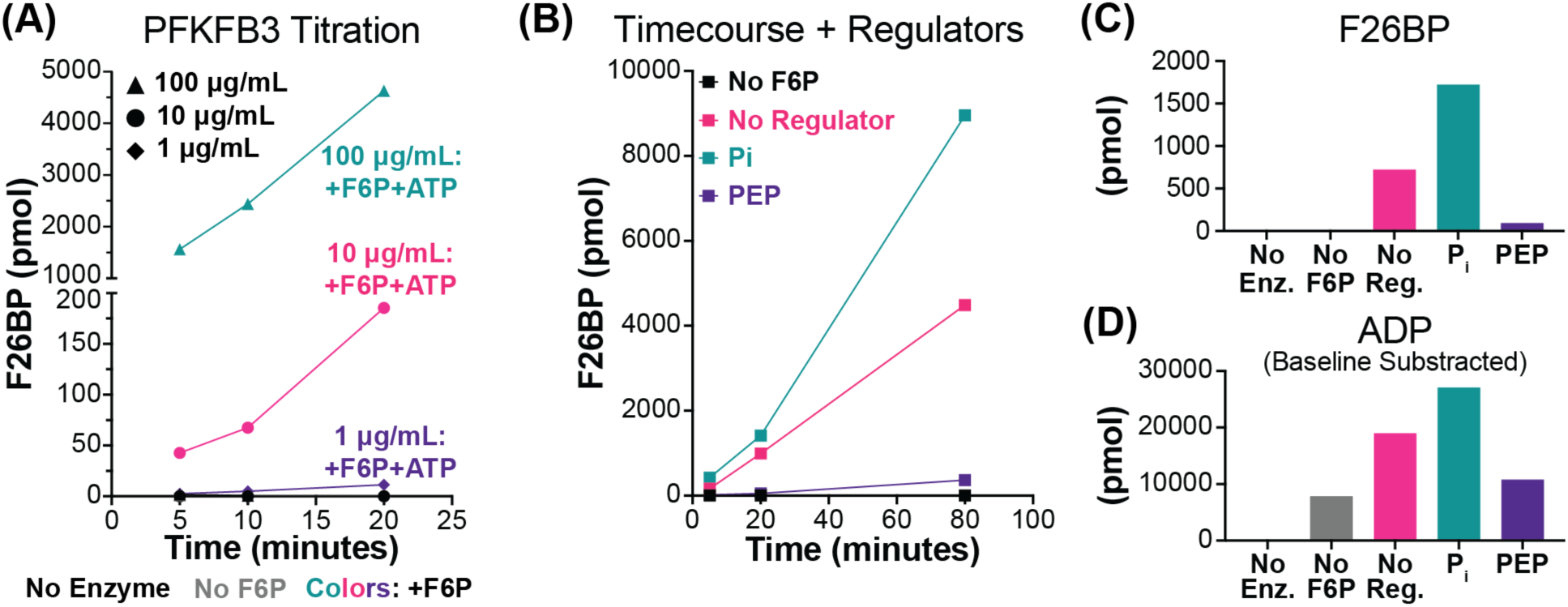
Validating recombinant PFKFB3 activity. (A) Time course with varying PFKFB3 titrations in the presence of 3 mM ATP and 0.2 mM F6P. 10 μg/mL (triangles), 10 μg/mL (circles), 1 μg/mL (diamonds), or 0 μg/mL (black, all shapes) PFKFB3 was incubated with (colors: teal - 100 μg/mL, pink - 10 μg/mL, purple - 1 μg/mL) or without F6P (grey, negative control). (B) PFKFB3 (40 μg/mL) time course in the presence or absence of known allosteric regulators. Reaction conditions included: no F6P (black, negative control), no regulator (pink), the addition of 1 mM P_i_ (teal, allosteric activator), and the addition of 1 mM PEP (purple, allosteric inhibitor). (C-D) Detection of PFKFB3 (100 μg/mL) kinase activity by incubating or 0 μg/mL (black; No Enz.) PFKFB3 with conditions described for 20 minutes. (C) Estimated F26BP content compared to (D) base-line subtracted ADP content. In cross-comparing the products of the PFKFB3 kinase domain, the F26BP and ADP quantities correlated across the different regulatory conditions.

